# Stage-Specific RNA Turnover Drives Small RNA Dynamics in *Arabidopsis* – *Colletotrichum* Interactions

**DOI:** 10.64898/2026.01.25.701602

**Authors:** Carolina E. Armijos, Thi-Thu-Huyen Chu, Richard J. O’ Connell, Blake C. Meyers, Patricia Baldrich

## Abstract

Small RNAs (sRNAs) are key regulators of plant defense and have been implicated in cross-kingdom interactions with pathogens. The hemibiotrophic fungus *Colletotrichum higginsianum* infects *Arabidopsis thaliana* through three stages: appressorial penetration, biotrophy, and necrotrophy. However, the dynamics of fungal and plant sRNA populations across these three stages have not been elucidated. Using high-throughput sequencing, we profiled sRNAs from *A. thaliana* and *C. higginsianum* during *in planta* appressorium (PA), biotrophic (BP), and necrotrophic (NP) phases, and compared them to fungal mycelia (MY) and *in vitro* appressoria (VA). Our analyses revealed stage-specific patterns in sRNA accumulation in both the plant and the pathogen. In *C. higginsianum*, sRNAs were dominated by 29 nt species in PA, BP, MY, and VA, but shifted to 18 nt in NP, consistent with RNA degradation during host cell death. In *A. thaliana*, sRNAs transitioned from 30-33-nt in PA/BP to a 21 nt dominant peak in NP. Also, TE-derived siRNAs and other regulatory sRNAs (miRNAs, ncRNA, snoRNAs and snRNAs) declined during NP. A total of 62 host miRNAs showed differential accumulation, including core plant developmental regulators active across infection stages, and stage-specific miRNAs such as miR396, miR170/171, miR472, and miR858b. tRFs displayed opposite trends in host and pathogen: fungal tRFs declined in NP, while host misc-tRFs, 5′-tRFs, and 3′-tRFs increased, suggesting contrasting regulatory roles. These results provide new insights into RNA-mediated plant-fungal interactions.

## 1. Introduction

*Colletotrichum higginsianum* is a hemibiotrophic ascomycete fungus responsible for causing anthracnose disease in various cruciferous plants (Damm et al. 2014; Yan et al. 2018). This includes species within the *Brassica* genus, such as *Brassica oleracea* (cabbage) and *Brassica rapa* (turnip), as well as *Raphanus* species like *Raphanus sativus* (radish) (Narusaka et al. 2006; Lee et al. 2018; Choi et al. 2019). The pathogen has become a significant threat to agriculture due to its potential to induce substantial economic losses (Yan et al. 2018). *C. higginsianum* employs a multistage hemibiotrophic infection process. First, a dome-shaped appressorium penetrates the host surface through a combination of mechanical pressure and localized enzymatic breakdown. Then, a bulbous biotrophic hyphae, which is enclosed by the host plasma membrane, grows within the living epidermal cells. Finally, the fungus transitions to a necrotrophic phase, where it produces thin and rapidly expanding hyphae that kill and degrade host tissues (O’Connell et al. 2012).

*C. higginsianum* also infects the model plant *Arabidopsis thaliana* (Tsushima et al., 2019). Since most *A. thaliana* ecotypes are susceptible to this fungus, it is considered well-adapted to infecting *A. thaliana* (Shimada et al. 2006). Consequently, its interaction with *Arabidopsis* serves as a valuable model system for studying the cellular and molecular mechanisms underlying fungal pathogenicity (Koch et al. 2025).

Plants have a sophisticated immune system. Pathogen-associated molecular pattern (PAMP)-triggered immunity (PTI), which detects conserved microbial molecules such as flagellin, chitin, and glycoproteins through membrane receptors called pattern recognition receptors (PRRs) and/or surface receptors like transmembrane receptor-like kinases (Jiang et al. 2023). Activation of PTI leads to the expression of pathogenesis-related (PR) genes, the generation of reactive oxygen species (ROS), callose deposition, and the accumulation of salicylic acid (SA) (David et al. 2019). As a response, pathogens secrete effector proteins into their host plants to suppress PTI, facilitating the infection (Zhang et al. 2022). Effector-triggered immunity (ETI) is another way plants detect pathogens through recognition of effectors directly and indirectly through resistance proteins. ETI triggers a stronger and faster immune response through resistance proteins known as nucleotide-binding-leucine-rich repeat-containing receptors (NLRs) (Nguyen et al. 2021).

By directly or indirectly regulating a wide variety of genes, small RNAs (sRNAs) are an intrinsic part of the plant defense responses (Jiang et al. 2023). They have even been suggested to function as antimicrobial agents by directly targeting pathogen genes through a process termed host-induced gene silencing (HIGS) (You et al. 2019). sRNAs are molecules of 18 to 30 nucleotides (nt), and function as crucial regulators of development, growth, reproduction, and biotic and abiotic stress responses (Zhan and Meyers 2023). They mainly fall into two major categories: microRNAs (miRNAs) and small interfering RNAs (siRNAs). miRNAs originate from endogenous MIR genes that get transcribed by RNA polymerase II (Pol II) into primary miRNAs (pri-miRNAs). Mature miRNAs are typically 20 to 22 nt long. siRNAs range from 21 to 24 nt in length and derive from long double-stranded RNA molecules (Baldrich et al. 2022; Bilir et al. 2022). Unlike miRNAs, siRNAs can originate from both endogenous genes and external sources such as viruses, transposons, and transgenes (Bilir et al. 2022).

This study aimed to identify and characterize sRNAs from the plant and the pathogen that may play roles in each phase of infection, namely *in planta*-formed appressoria, biotrophic and necrotrophic phases, by comparing their profiles with those in axenically grown fungal mycelia and *in* vitro-formed appressoria. To that end, we sequenced sRNAs and did comparative accumulation analysis of different types of sRNAs, between each phase of the infection cycle. Identifying these sRNAs could facilitate the development of sustainable strategies for controlling fungal infections, with potential applications not only in *A. thaliana* but also in other plant species.

## 2. Materials and Methods

### Preparation of plant and fungal materials

*C. higginsianum* strain IMI 349063A was cultured for spore production on Mathur’s agar medium, as described previously (Rutter et al. 2022). For the mass production of fungal appressoria *in vitro*, 40 ml of spore suspension (2 x 10^6^ spores/ml) was poured into a 12-cm-square polystyrene Petri dish and the spores were allowed to settle for 40 min. After covering the base of the dish with a 12-cm-square piece of nylon mesh (50 μm pore size), excess water was decanted and the plate was incubated for 22 hpi at 25°C in a humid chamber. The nylon mesh was then gently floated off with water and residual liquid was removed by vigorous shaking before adding 25 ml of TRIzol reagent (ThermoFisher Scientific) and disrupting the appressoria using a cell scraper (Corning, product no. 3011). Total RNA isolation was performed according to the manufacturer’s instructions. Each biological replicate comprised pooled extracts from five Petri dishes (approximately 4 x 10^8^ appressoria in total).

To obtain undifferentiated fungal mycelia, potato dextrose broth (100 ml) in a 250 ml Erlenmeyer flask was inoculated with spore suspension to a final concentration of 1 x 10^6^ spores/ml and shaken (100 rpm) at 25°C. After 3 days, mycelium was harvested by filtration through a nylon mesh before grinding in liquid nitrogen in a pre-cooled mortar and pestle and total RNA was extracted with TRIzol.

For harvesting RNA from infected leaves, plants of *A. thaliana* ecotype Col-0 were grown in a peat-based compost (Floradur-B, Floragard, Oldenburg, Germany) using a Percival AR-36L3 growth chamber (CLF PlantClimatics GmbH, Wertingen, Germany; 12-h photoperiod, 230 μmol m^-2^ s^-1^ photon flux density, 23/21°C day/night temperature, 70% relative humidity). Fully-expanded rosette leaves were excised from 5-week-old plants and the abaxial surface was brush-inoculated with approximately 100 μl spore suspension (5 x 10^6^ spores/ml), using pieces of nylon mesh (50 μm pore size, 1 cm x 2cm) to maintain a thin film of inoculum over the entire leaf surface. Inoculated leaves were incubated in humid boxes in the dark at 25 °C.

*In planta* appressoria (PA), comprising pre-penetration appressoria on the leaf surface, and the early biotrophic phase (BP), comprising post-penetration appressoria and young biotrophic hyphae, were harvested by epidermal stripping at 22 hpi and 40 hpi, respectively. For this, the adaxial leaf surface was first adhered to a Petri dish with double-sided tape and then pieces of infected epidermis were peeled off using tweezers and immediately flash-frozen in liquid nitrogen. For sampling the necrotrophic phase (NP), areas of necrotic leaf tissue that appeared transparent and water-soaked at 60 hpi were excised using a scalpel and flash-frozen in liquid nitrogen. All samples were then ground in liquid nitrogen and total RNA was extracted as described above. Each biological replicate comprised tissue harvested from approximately 30 leaves per time-point, per condition.

The quality and quantity of all extracted total RNA samples was measured by Qubit fluorometer assay and by running RNA denaturing agarose gels. Prior to preparing sRNA libraries, samples were dried in GenTegraRNA tubes (GenTegra, Pleasanton, CA 94566, USA) and stored at ambient temperature.

### Annotation of A. thaliana and C. higginsianum genome

The A. *thaliana* genome assembly (TAIR10) and annotation file were downloaded from TAIR database (https://www.arabidopsis.org). The *C. higginsianum* genome assembly and annotation file were downloaded from the JGI Genome Portal (https://genome.jgi.doe.gov/portal/).

For the *C. higginsianum* genome, repetitive regions were annotated with RepeatMasker v4.1.5 (Smit et al. 2015). A RepeatModeler database was first created from the genome assembly using the *BuildDatabase* command with default parameters. RepeatModeler v2.0.3 (Smit and Hubley 2015) was then run on the genome assembly using the *- LTRStructoption* flag to enable identification of Long Terminal Repeats (LTR) retrotransposon structures. Two rounds of classification refinement were performed using RepClassifier GPLv2 (Smit and Hubley 2015). Classified and unclassified repeats were combined with the RepBase v23.08 *fngrep* reference database into a comprehensive repeat library. Finally, repeats were masked in the genome assembly using RepeatMasker. rRNA genes were predicted by RNAMMER (Lagesen et al. 2007) using default parameters. tRNAs genes were predicted by tRNAscan-SE 2.0 (Chan et al. 2021) using default parameters. For *A. thaliana* and *C. higginsianum* genomes, tRNA-derived fragments were annotated using Unitas v1.8.0 (Gebert et al. 2017) with default parameters.

### Library preparation, sequencing, and analysis

sRNA libraries were constructed using NEBNext^®^ Small RNA Library Prep Set for Illumina^®^ following the manufacturer’s instructions. Libraries were combined and sequenced with the Illumina NextSeq 500 platform in single-end mode (100 bp) at the University of Delaware DNA Sequencing & Genotyping Center (Newark, DE, USA). Sequencing adapters were trimmed using Cutadapt v4.10 (Martin 2011) with the following parameters: *-a TGGAATTCTCGGGTGCCAAGGAACTCCAGTCAC-g GTTCAGAGTTCTACAGTCCGACGATC-u 1-m 10-j 0 --max-n 0-o sample.fq*. Read quality was assessed using MultiQC v1.8 (Ewels et al. 2016). Clean reads were aligned to *A. thaliana* and *C. higginsianum* genomes and their corresponding genomic features using Bowtie v2.4.5 (Langmead and Salzberg 2012). The raw read counts were normalized, and the resulting data were visualized through plots generated using R v4.3.0 (R Core Team 2023).

miRNAs for *A. thaliana* were identified using the most recent version of miRBase (https://www.mirbase.org), sequences specific to the *Arabidopsis thaliana*. Differential accumulation (DA) analysis of miRNAs was performed using DESeq2 v1.40.2 (Love et al. 2014) in R. Input data consisted of raw miRNA read counts derived from Bowtie2 alignments. Wald tests were used to compute p-values for differential expression between conditions with a significance threshold of α = 0.05. Plots were generated using *ggplot2* package v3.4.4 (Wickham 2016) for exploratory data analysis and interpretation. The raw read counts from the Unitas pipeline were normalized, and visualizations were also created using R.

## 3. Results

### sRNA profiles are distinct between different stages of the *C. higginsianum* infection cycle

To identify the sRNAs involved in the infection process of *A. thaliana* by *C. higginsianum*, we generated sRNA libraries from five distinct stages, each with three biological replicates. Three infection stages, namely *in planta*-formed appressoria (PA), biotrophic (BP), and necrotrophic (NP) phases, were compared with two axenic fungal stages, namely liquid-cultured mycelia (MY) and *in vitro*-formed appressoria (VA). To identify sRNAs originating from both the plant and fungal sides, we mapped all reads to the respective reference genomes: TAIR10 for *A. thaliana* (https://www.arabidopsis.org) and Colhig2 for *C. higginsianum* (https://genome.jgi.doe.gov/portal/). Approximately 70 to 80% of sRNA reads mapped to the *A. thaliana* genome during the PA, BP, and NP phases, while 5 to 17% mapped the *C. higginsianum* genome. This is consistent with these stages representing active interaction between *C. higginsianum* and *A. thaliana*, where *Arabidopsis* accounts for most of the biomass used for RNA extraction. In contrast, during the MY and VA phases, 40 to 70% of the reads mapped to the *C. higginsianum* genome (Figure S1).

To further characterize the sRNA populations, we analyzed the size distribution of sRNAs across the different infection (PA, BP, NP) and control (MY and VA) stages. In alignment with typical fungal sRNA classes (He et al. 2023), *C. higginsianum* sRNAs displayed a prominent peak at 29 nt across all stages except for the NP phase (Figure S2A). The NP displayed a markedly different size distribution, characterized by a peak at 18 nt. On the other hand, *A. thaliana* sRNAs in PA and BP both exhibit their highest peaks at 30 nt, whereas the NP phase shows its maximum peak at 21 nt. In both PA and BP, secondary peaks occur at 18 nt, 21 nt, 24 nt, and 33 nt. Meanwhile, the NP phase displays high abundance levels across a broad range from 18 nt to 34 nt (Figure S2B). This illustrates the differences in sRNA populations, not only between organisms, but also between the different stages of the infection process.

### Infection progression is marked by a decline in host and pathogen sRNAs

To understand the genomic origin of sRNAs from the infection process, we annotated common features from *C. higginsianum* genome, including ribosomal RNAs (rRNAs), transfer RNAs (tRNAs), and repetitive elements, given that this information was not available for this species. For rRNAs, we obtained 223 genes corresponding to 145 8S, 40 18S, and 38 28S (Table S1). We also annotated 354 tRNAs (Table S2). We further found that 5.53% of the genome consisted of repetitive elements, where 2.00% were retroelements and 2.58% were DNA transposons, 0.01% were rolling circles, and 0.96% were unclassified (Table S3). These annotations provide a foundational genomic context for investigating the origin and potential function of *C. higginsianum*-derived sRNAs during *C. higginsianum* infection of *A. thaliana*.

To investigate how sRNAs are distributed across genomic features during *C. higginsianum* infection, we analyzed sRNA abundance at PA, BP, NP, MY, and VA stages and categorized the reads by genome feature. We observed variation in sRNA abundance across different infection stages of *C. higginsianum*, categorized by genome features (Figure 1A). Among all features, TEs and rRNAs exhibit the highest sRNA abundance. These two features display similar expression patterns across the five infection phases, with TEs consistently showing greater abundance than rRNAs. In contrast, genes and CDSs show moderately lower sRNA abundance compared to TEs and rRNAs. Their levels peak during the MY and VA stages, decline during the PA and BP stages, and drop substantially in the NP phase. tRNAs exhibit the lowest sRNA abundance among the genomic features analyzed. Their levels are highest during the PA and BP phases, slightly decline during the MY and VA phases (particularly in VA), and show a sharp decrease in the NP phase. All these suggest that sRNA levels in *C. higginsianum* vary by feature and stage, dropping notably in the NP phase.

**Figure 1.**
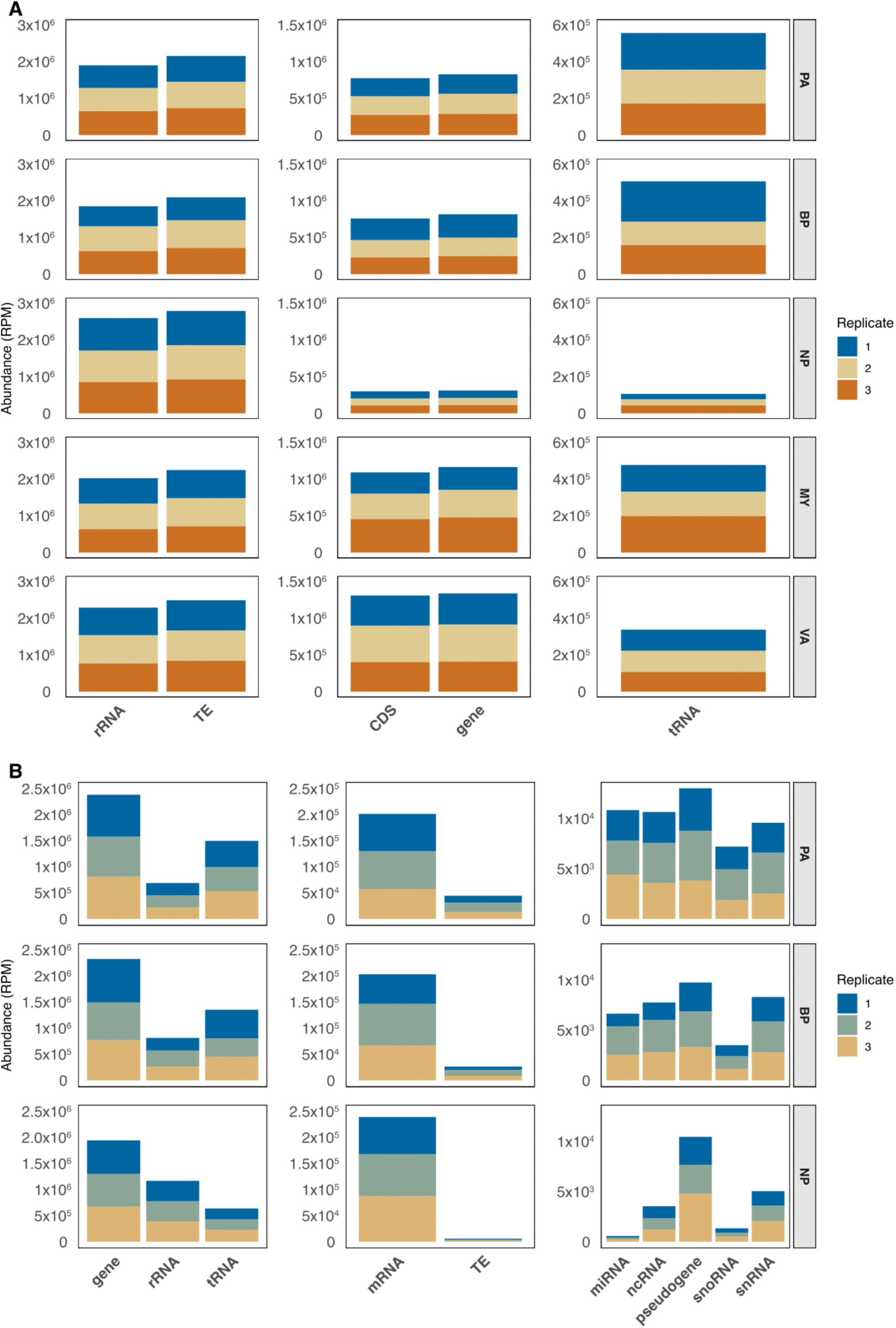
Stage-specific gene expression dynamics in *C. higginsianum* and *A. thaliana* during infection. sRNA mapping to each genomic feature for *C. higginsianum* (B) and *A. thaliana* (A). A) The y-axis displays RNA abundance in reads per million (RPM) for the different stages of *C. higginsianum*: *In Planta* Appressorium (PA), Biotrophic Phase (BP), Necrotrophic Phase (NP), *In Vitro* Appressorium (VA), and Mycelia (MY). Three replicates are indicated by distinct colors (blue, mustard-yellow, and orange). The x-axis represents various *C. higginsianum* features, including ribosomal rRNAs (rRNA), Transposable Elements (TE), coding DNA Sequences (CDS), genes, and transfer RNAs (tRNAs). B) The RNA abundance (RPM) in *A. thaliana* during the PA, BP, and NP infection stages is shown on the y-axis. The *x-axis* illustrates the corresponding features of *A. thaliana*, such as genes, rRNA, tRNA, messenger RNAs (mRNAs), micro RNAs (miRNAs), non-coding RNAs (ncRNAs), pseudogenes, small nucleolar RNAs (snoRNAs), and small nuclear RNAs (snRNAs). Three replicates are represented using different colors (blue, green, and mustard-yellow).

On the *A. thaliana* side, genes exhibit the highest sRNA abundance across all stages, with relatively stable levels during the PA and BP phases, followed by a slight decrease in the NP phase (Figure 1B). rRNAs are the second most abundant feature, with a modest increase in the NP phase compared to PA and BP. tRNAs follow, showing higher sRNA levels in PA and BP, which decline in NP. TEs display a marked decrease in sRNA abundance from PA to NP. A similar downward trend is observed for miRNAs, ncRNAs, snoRNAs, and snRNAs. This suggests that sRNA abundance across each *A. thaliana* genome feature (except genes) progressively declines during infection, mirroring the trend observed in *C. higginsianum*.

### Size distribution of C. higginsianum sRNA fragments indicates a higher RNA turnover during the NP phase

Analyzing the size distribution of reads mapped to each feature provides insights into their processing and turnover across infection stages. The size distribution profiles of *C. higginsianum* sRNAs mapped to rRNA sequences revealed a predominant peak at 29 nt in the PA, BP, MY, and VA stages. In contrast, the NP stage exhibited a primary peak at 18 nt, accompanied by a broad distribution of sRNA sizes ranging from 19 to 32 nt. While the MY and VA stages were characterized solely by the dominant 29 nt peak, the PA and BP stages showed an additional secondary peak at 32 nt, which increased in abundance from PA to BP. In the NP stage, both the 29 nt and 32 nt peaks remained detectable but were no longer the most abundant features (Figure S3). We hypothesize that the differences observed in rRNA fragment profiles may correspond to senescence processes and cell death occurring at the necrotrophic stage. We observed a similar pattern in the size distribution profiles for *C. higginsianum* sRNA reads that mapped to TEs (Figure S4). There is a predominant peak at 29 nt among the TE-derived RNAs during the PA, BP, MY, and VA stages, whereas in the NP stage, the dominant peak shifts to 18 nt, accompanied by a broad distribution of sRNA sizes ranging from 19 to 32 nt. This shift in the NP stage suggests an increased breakdown of TE-derived RNAs, in line with the overall higher RNA turnover observed during necrotrophy.

Reads mapping to fungal gene features exhibited a somewhat analogous pattern to plants of size distribution profiles; however, in this case, we observed a single notable peak at 29 nt across the PA, BP, MY, and VA phases. In the NP phase, the predominant peak was at 18 nt, followed by a smaller peak at 29 nt (Figure S5). This reinforces our hypothesis that the NP phase has a higher degradation pattern of smaller reads, indicating a higher RNA turnover, possibly due to cell death induced during the NP.

Finally, the size distribution profiles of tRNA-derived fragments displayed distinct patterns compared to other RNA features. The dominant peaks varied slightly between stages: in order of abundance, 35 nt and 36 nt in PA; 35 nt and 34 nt in BP; 34 nt and 35 nt in NP; and consistently 34 nt and 35 nt in both VA and MY. Notably, the VA and MY stages exhibited the lowest overall abundance of tRNA-derived fragments across all samples (Figure S6). This reduced abundance may reflect lower tRNA cleavage activity or decreased tRNA turnover during these non-plant stages, possibly linked to reduced translational demand or altered stress responses within the fungal structures, in the absence of the host plant.

### Size distribution of *A. thaliana* sRNA fragments also indicates a higher RNA turnover during the NP phase

We did the same fragment size distribution analysis as previously described, using the *A. thaliana* features annotated in the TAIR10 genome (https://www.arabidopsis.org). Size distribution patterns for gene-and rRNA-derived reads showed a similar trend. In both features, PA and BP stages exhibited prominent peaks at 30 - 33 nt, followed by peaks at 21 nt. In contrast, the NP phase was characterized by a shift toward shorter fragments, with the dominant peak at 21 nt and a broad distribution of reads from 18 - 32 nt (Figures S7 and S8). These changes suggest enhanced RNA turnover during the NP phase, consistent with the patterns observed in *C. higginsianum*. Similarly, ncRNA-derived reads showed a major peak at 21 nt in both PA and BP. In the NP phase, this 21 nt peak was maintained but accompanied by higher abundance across a broader range of 18–24 nt (Figure S9). This also reflects increased degradation activity during NP.

In contrast, size distribution profiles for TE-, miRNA-, and snRNA-derived reads all declined sharply in NP. TE-derived reads displayed strong 23 - 24 nt peaks in PA and BP, but these peaks were lost in NP, with an overall reduction in mapped reads (Figure S10). Similarly, miRNAs showed a consistent 21 nt peak across all stages, but their abundance dropped substantially from PA to NP (Figure S11). snRNAs followed the same trend: while PA and BP had clear peaks at 24 - 26 nt, the NP phase exhibited reduced overall accumulation and a flatter distribution (Figure S12). Together, these results indicate a broad loss of sRNA accumulation from these features during late infection, likely reflecting degradation dynamics.

tRNA-derived sRNAs exhibited a distinct pattern. The BP phase showed the highest accumulation, with prominent peaks at 31, 30, and 32 nt. In NP, the peak shifted to 30 nt with lower adjacent peaks at 29 and 31 nt. The PA phase differed entirely, showing broader peaks spanning 29 - 35 nt but at much lower abundance overall (Figure S13). This indicates stage-specific processing of tRNA-derived sRNAs, with reduced accumulation in PA.

Finally, snoRNA-derived reads remained consistently low and relatively stable across all stages. The PA and BP phases displayed small peaks at 21 - 23 nt, while NP was dominated by a modest 21 nt peak. Unlike other features, snoRNAs did not exhibit a pronounced decline in NP (Figure S14).

### A core of differentially accumulated *A. thaliana* miRNAs is present throughout all stages of infection

To understand the role of *A. thaliana* miRNAs in each of the phases of infection, we did a quantification and differential accumulation analysis of known miRNAs in the three different phases. We observed dynamic changes in miRNA expression in *A. thaliana* during different infection stages of *C. higginsianum*, with distinct regulatory patterns observed across comparisons. Out of the 350 miRNAs analyzed, 62 miRNAs showed a differential accumulation in at least one comparison. We found a core of 26 miRNAs that are differentially accumulated in all three comparisons.

To better understand which miRNAs were up-or down-regulated during infection, we separated them into these two categories and analyzed how many were shared across stage comparisons. Our quantitative analysis of upregulated miRNAs (log2FC > 0) revealed that 44 miRNAs were upregulated in the three comparisons. Thirteen miRNAs were upregulated in BP vs. PA and NP vs. PA. Only two miRNAs were upregulated, both in BP vs PA and NP vs. BP (Figure 2B). For instance, miR396b-5p, miR159a, miR159-3p were consistently significantly up-regulated across all comparisons. miR398bc-5p was also upregulated in all phases, but only NP/BP vs PA phase comparisons were significant. miRNA858a was also upregulated in all phases, but only NP vs PA phase comparison was significant. This suggests that these miRNAs play a steady role during the whole infection process.

**Figure 2.**
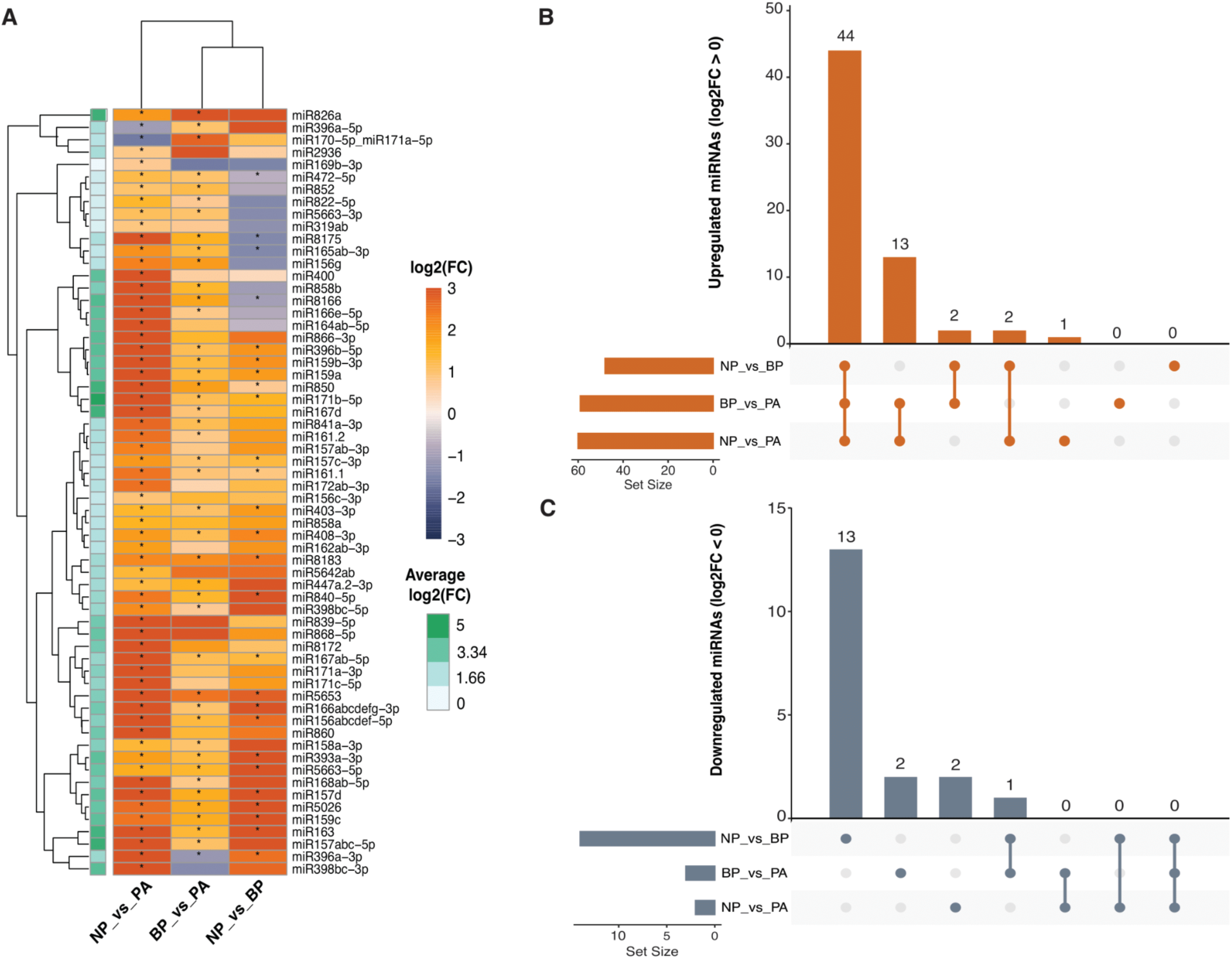
Differential accumulation (DA) of miRNAs in *A. thaliana* during *C. higginsianum* infection suggests that miRNAs profiles are most distinct between the PA and NP. A) Heatmap showing log2 fold changes (log2FC) of miRNAs across of *C. higginsianum* stages: *In Planta* Appressorium (PA) vs. Biotrophic Phase (BP), BP vs. Necrotrophic Phase (NP), and NP vs. PA. Dark orange and lavender indicate up-and downregulated miRNAs, respectively. B and C) Upset plots summarizing the number of significantly B) downregulated (log2FC < 0) and C) upregulated (log2FC > 0) miRNAs in each *C. higginsianum* stage.

In contrast, downregulated miRNAs (log2FC < 0) had a limited overlap and a narrower distribution (Figure 2C). Only two miRNAs were downregulated, both in BP vs. PA and NP vs. PA. Thirteen miRNAs were downregulated in NP vs. BP. We did not observe shared downregulated miRNAs across all three phase comparisons, but we did observe one miRNA (miR169b-3p) that was downregulated both in BP vs. PA and NP vs. BP. These results suggest that miRNA up-regulation is more prominent and stage-specific, while down-regulation is relatively limited during infection.

### Few *A. thaliana* miRNAs show stage-specific patterns during fungal infection

Additionally, 17 miRNAs had a distinct differential accumulation pattern in the different comparisons, suggesting a significant phase-specific role. For example, miR396a-5p and miR170-5p_miR171a-5p were significantly down-regulated in NP compared to PA (NP vs. PA) but significantly upregulated in BP vs. PA. In the NP vs. BP phase comparison, miR396a-5p and miR170-5p_miR171a-5p were also upregulated, but only miR170-5p_miR171a-5p expression was significant. This suggests that these miRNAs play a role specifically during the BP stage. Additionally, miR472-5p and miR165ab-3p showed contrasting expression trends; they were significantly down-regulated in NP compared to the BP but significantly up-regulated in NP vs. PA and BP vs. PA. Similarly, miR858b was significantly downregulated in the NP phase compared to the BP but non-significantly upregulated in NP vs. PA and BP vs. PA, suggesting a role in fungal early/pre-penetration and the NP phase. Notably, miR396a-3p was significantly downregulated during the BP phase compared to the PA, while it was significantly upregulated in NP vs. PA and NP vs. BP, suggesting distinct roles in biotrophic establishment and necrotrophic transition. Conversely, miR398bc-3p was downregulated during the BP phase compared to the PA, although this change was not statistically significant. It was also significantly upregulated in NP vs. PA but not significantly upregulated in NP vs. BP. miR169b-3p exhibited a non-significant downregulation in BP vs. PA and NP vs. BP, but was significantly upregulated in NP vs. PA, collectively implicating late infection-specific roles.

### tRF abundance varies by infection stage both in *A. thaliana* and *C. higginsianum*

To investigate tRFs dynamics in the plant-fungus interaction, we analyzed their abundance across different infection stages using Unitas software (Gebert et al. 2017). The identified tRFs come from a classification based on their cleavage position in the mature tRNA and pre-tRNA structure (Figure 3A and 3B). These include tRFs derived from pre-tRNA, such as the 5’ leader sequence (tRNA-leader) and the 3’ trailer sequence containing a poly-U terminus (tRF-1s), and fragments from the mature tRNA body, like 5’-tRFs, 3’-tRFs (with and without the CCA tail), both 5’/3’-tR-halves, and miscellaneous tRFs (misc-tRFs) that map internally without aligning to a defined terminal region (Akiyama and Ivanov 2023). We observed distinct patterns of tRF abundance during *C. higginsianum* infection in *A. thaliana* (Figure 3C). Among all tRF types, 3′-tRNA halves were the most abundant across all infection phases. Misc-tRFs ranked next, showing high levels in PA and BP but dropping to lower, stable levels in NP, MY, and VA. 5′-tRNA halves displayed a similar trend: abundant in PA/BP, sharply reduced in NP, MY, and VA. 3′-CCA-tRFs were less abundant overall but showed a distinct peak in the MY phase, while maintaining low levels during PA, BP, NP, and VA. 5′-tRFs followed, with relatively stable levels in PA and BP that decreased in NP, MY, and VA. Next in abundance were 3′-tRFs, which showed their highest levels in the MY phase, moderate levels in PA, BP, and VA, and the lowest levels in NP. Finally, tRF-1 and tRNA leader fragments had the lowest abundances among all tRF types. Both showed a peak in BP and remaining low otherwise (Figure 3C). Overall, tRF levels shifted dynamically, with many types reduced during NP.

**Figure 3.**
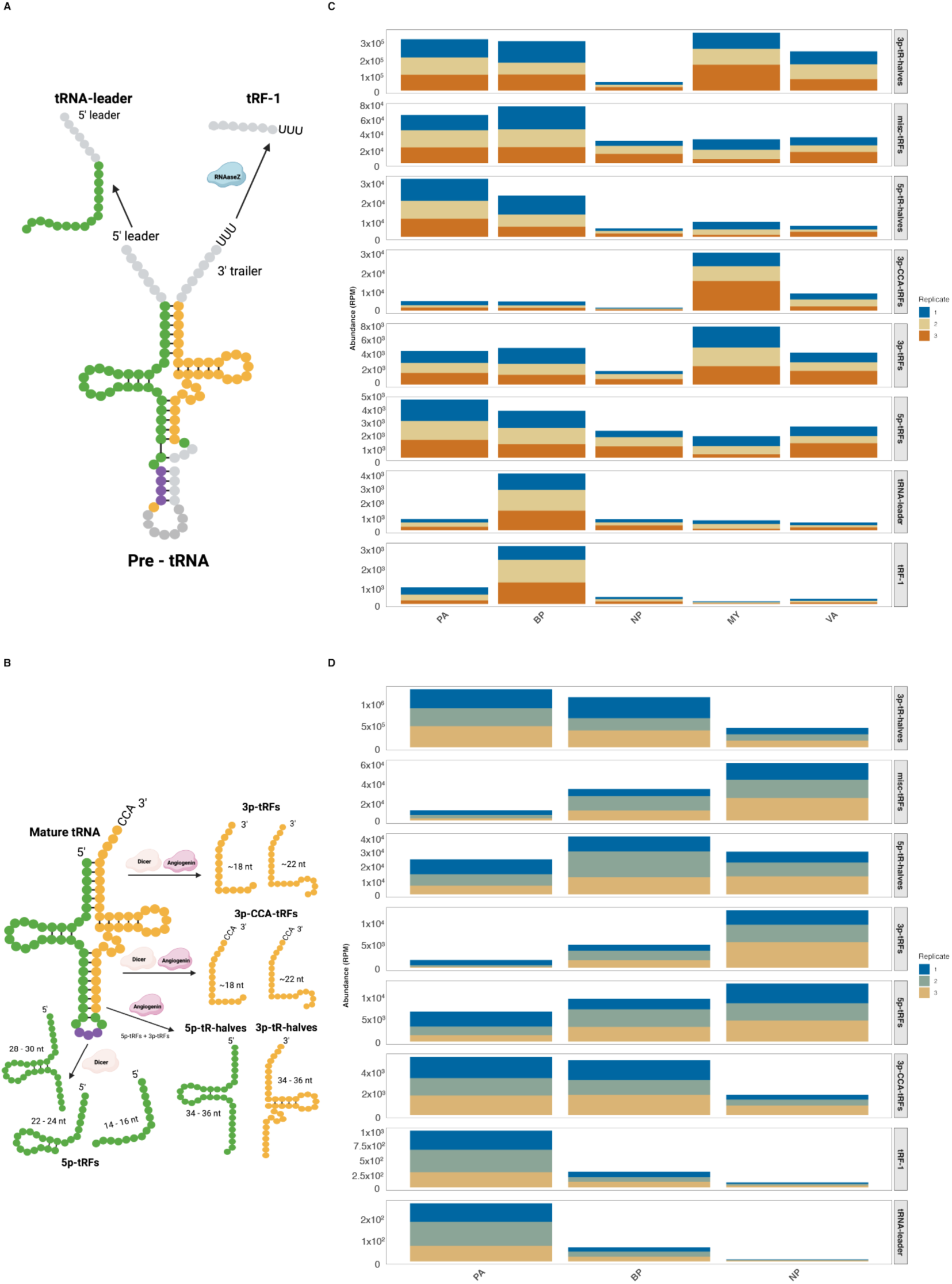
Abundance of tRNA-derived fragments (tRFs) during *C. higginsianum* infection in *A. thaliana* is stage specific. A) and B) Schematic representation of various types of tRFs categorized by their origin within the tRNA structure. tRNA-leaders derive from the pre-tRNA structure 5’-leader sequence. tRF-1s are tiny segments that originate from the 3′ ends of pre-tRNA, which include the’poly-U’ sequence at their 3′ terminus. 5′ tRFs are derived from the 5′ end up to the D-loop; 5′ tR-halves extend from the 5′ end to the anticodon loop; 3′ tRFs span from the TψC-loop to the 3′ end (excluding the CCA tail); 3′ CCA-tRFs originate from the TψC-loop to the 3′ CCA sequence; 3′ tR-halves extend from the anticodon loop to the 3′ end; and miscellaneous tRFs (misc-tRFs) are those that map to mature tRNA without aligning precisely to the defined terminal regions (Akiyama and Ivanov 2023). C) The y-axis displays RNA abundance in reads per million (RPM) for the different stages of *C. higginsianum*: *In Planta* Appressorium (PA), Biotrophic Phase (BP), Necrotrophic Phase (NP), *In Vitro* Appressorium (VA), and Mycelia (MY). Three replicates are indicated by distinct colors (blue, mustard-yellow, and orange). The x-axis represents different tRF types. D) The RNA abundance (RPM) in *A. thaliana* during the PA, BP, and NP infection stages is shown on the y-axis. The x-axis represents different tRF types. Three replicates are represented using different colors (blue, green, and mustard-yellow).

We observed a similar phenomenon in *A. thaliana* (Figure 3D). The most abundant tRF types mirrored those in *C. higginsianum*, and each type showed stage-specific fluctuations, with some peaking in BP and others declining by NP. When comparing tRNA abundances between *C. higginsianum* and *A. thaliana* across the PA, BP, and NP phases, we observe that the abundance of 3p-tRNA halves is higher in *A. thaliana* than in *C. higginsianum,* exhibiting a decreasing pattern of abundance in both species from PA to NP. In *C. higginsianum*, the abundance of mic-tRFs decreased across the phases, with counts dropping substantially in NP. Conversely, *A. thaliana* exhibits an increase in mic-tRFs, rising from PA to in NP. For 5p-tRNA halves, we observed a significant difference in the BP phase, where *A. thaliana* demonstrated a notably higher abundance compared to *C. higginsianum*. Both species exhibit similar low abundance levels for 3p-CCA-tRFs. In *C. higginsianum*, 5p-tRFs decreased in the PA phase to the NP phase, while in *A. thaliana*, they remained steady in the PA phase but decreased in the NP phase. For 3p-tRFs, in *C. higginsianum* they showed a decline from PA to NP, whereas in *A. thaliana* they increased from PA to NP. For tRF-1, in *C. higginsianum* they peak during the BP phase, while *A. thaliana* they reach their peak in the PA phase. Lastly, tRNA leader abundance remained relatively low across all phases in *A. thaliana*, while in *C. higginsianum* there was a higher abundance in the PB phase. These results highlight species-specific differences in tRF dynamics, suggesting distinct regulatory roles for tRNA fragments in the host and the pathogen during infection.

## 4. Discussion

### Distinct sRNA profiles reflect stage-specific regulatory dynamics

The evaluation of read length distributions is a powerful tool for inferring the role of different sRNAs in any given process (Silvestri et al. 2019). In *C. higginsianum*, the dominant 29 nt sRNA peak during the PA, BP, MY, and VA phases aligns with non-canonical RNAi pathways, particularly Dicer-independent mechanisms involving RNA-dependent RNA polymerase (RdRP) (Weiberg et al. 2013; Wang et al. 2016; 2022). These sRNAs are often linked to fungal virulence, as observed in other pathogens like *Botrytis cinerea*. In contrast, the NP phase exhibits a distinct 18 nt peak and a broad size distribution, suggesting either an increased degradation of fungal transcripts by the host, a stress response in the pathogen, or increased RNA degradation during autophagy/senescence of older hyphal compartments (Melnyk et al. 2011). The sharp decline in TE- and rRNA-derived sRNAs during NP further supports a breakdown in fungal RNAi-mediated defense response, likely due to host counterattacks or necrotrophic transition-associated RNA turnover.

In *A. thaliana*, the PA/BP phases are marked by 30 - 33 nt sRNAs (putatively DCL-independent) and secondary peaks at 21 - 24 nt (canonical DCL products), indicative of active RNA-directed DNA methylation (RdDM) and transcriptional silencing (Melnyk et al. 2011). The shift to a dominant 21 nt peak in NP suggests a transition to post-transcriptional gene silencing (Cao et al. 2014), while the broad smears (18 - 34 nt) imply widespread RNA degradation or sRNA populations (e.g. tRFs). The decline in TE-associated 24-nt siRNAs and other regulatory sRNAs (miRNAs, ncRNAs) by NP highlights the plant’s diminishing capacity to regulate immune responses as the infection progresses.

### Distinct sRNA feature sources also reflect stage-specific host-pathogen interactions

During the infection process, *C. higginsianum* and *A. thaliana* exhibit distinct but coordinated changes in the origin and abundance of sRNAs across different stages. In *C. higginsianum*, TEs and rRNAs constitute the main sources of sRNAs, with consistently high levels observed during PA and BP phases, followed by a pronounced reduction in the NP. Although tRNA fragments contribute less overall, they display a notable increase in sRNA production during the PA/BP stages, indicating possible stress-induced, stage-specific cleavage. This accumulation of specific tRNA-derived fragments might be functionally significant, as they are emerging as key signaling molecules in cross-kingdom communication between plants and pathogens (Kusch et al. 2023).

In contrast, *A. thaliana* primarily generates sRNAs from genes, with stable levels during PA/BP that decline in NP. Additional sources, such as TEs, miRNAs, ncRNAs, snoRNAs, and snRNAs, show a similar decreasing trend from PA to NP, aligning with the fungal profile. These observations suggest that the two organisms exhibit different sRNA origins but a synchronized temporal pattern, suggesting interlinked regulation or shared trigger processes during infection progression.

### Some miRNAs show differential accumulation across all infection stages in A. thaliana’s battle against C. higginsianum

Our study reveals a complex miRNA dynamic in the host during infection, with some miRNAs playing a role during all different infection phases. For instance, miR393 demonstrated increased abundance in *Arabidopsis* infected with *Pseudomonas syringe* pv. tomato (Zhang et al. 2021) and cassava infected with *C. higginsianum* (Pinweha et al. 2015). Our results show that miR393a-3p is upregulated in all three phases, which is consistent with the upregulation of miR393 reported by Pinweha et al. (2015) and suggests that sustained suppression of auxin signaling (as it targets *TIR1*, a key auxin receptor) is a conserved response among different plant pathogen interactions.

Furthermore, the miR159-GAMYB pathway is highly conserved across various plant species, including *Arabidopsis*, tobacco, and rice (de Oliveira Cabral et al. 2024; Zheng et al. 2020). GAMYB is a transcription factor within the MYB family that regulates gene expression involved in plant growth and development, particularly in response to gibberellin hormone signaling (Liu et al. 2024). In tobacco, inhibition of miR159 results in GAMYB-driven activation that triggers Pathogenesis-Related (PR) genes (Zheng et al. 2020). In our study, the upregulation of miR159a and miR159b-3p across all infection stages suggests that this pathway remains active during *C. higginsianum* infection. However, in *A. thaliana*, miR159 may function to repress GAMYB during infection, potentially preventing deleterious effects such as stunted growth or programmed cell death, which could help avoid even more self-inflicted damage and provide long-term resistance without developmental costs. This could imply that miR159 has a context-and species-dependent role.

miR396b-5p is also upregulated across all comparisons. According to our results, miR396a-5p downregulation in NP vs. PA aligns with the findings of Soto-Suárez et al. (2017) findings, as they found that lower miR396a-5p would release Growth Regulating Factor (GRF) transcriptional factors to amplify necrotrophic defenses in plants like *A. thaliana*.

Finally, according to our results, miR858a is significantly upregulated across all three infection phases (PA, BP, and NP). This consistent high expression is notable because prior research by Camargo-Ramírez et al. (2018) identified miR858 as a negative regulator of disease resistance in *Arabidopsis* during the infection of *C. higginsianum*. As a result, its overexpression increased susceptibility to the fungal pathogen. Therefore, the sustained upregulation of miR858a suggests a suppression or delay of the plant’s defense response throughout the infection process.

### Other miRNAs show phase-specific regulation in *A. thaliana*’s infection with *C. higginsianum*

Our results revealed that different miR396 family members display opposing regulatory roles: miR396a-3p is downregulated in BP vs. PA but upregulated in NP vs. PA and NP vs. BP. It was demonstrated that miR396 negatively regulates immunity against fungal pathogens by targeting Growth Regulating Factor (GRF) transcriptional factors (Soto-Suárez et al. 2017). In fact, these authors showed that reduced levels of miR396 in *A. thaliana* MIM396 plants enhance resistance to fungal pathogens such as *P. cucumerina* and *C. higginsianum*, while miR396 overexpression increases susceptibility. Hence, during infection, miR396 levels gradually decline, enabling GRFs to activate defense responses like Reactive Oxygen Species (ROS), callose deposition, and Jasmonic Acid (JA) and ethylene (ET) signaling.

The miR170/miR171 are known to regulate Scarecrow-like transcriptional factors in Arabidopsis (Bologna et al. 2013). Among those, Scarecrow-like 14 (SCL14), which is a regulatory protein member of the GRAS family, is involved in activating genes related to detoxifying harmful compounds, such as xenobiotics and other toxic metabolites (Fode et al. 2008). Our results show that miR170-5p and miR171a-5p are downregulated in the NP vs. PA stage but upregulated in both the BP vs. PA and NP vs. BP phases. Therefore, it is plausible that miR170-5p and miR171a-5p may play a defensive role via the SCL transcription factor family to help the plant respond to metabolites produced by the pathogen during the necrotrophic phase of the infection.

The expression of miR858b in this study appears to be highly dependent on the infection phase. The overexpression of miR858 increases susceptibility to fungal pathogens, whereas its inhibition enhances resistance by promoting the upregulation of flavonoid-specific MYB transcription factors and the accumulation of antifungal compounds such as kaempferol and other phenylpropanoids (Camargo-Ramírez et al. 2018). In our results, miR858b was upregulated in NP vs. PA and BP vs. PA, but downregulated in NP vs. BP. This pattern suggests that the downregulation of miR858b in the NP relative to the BP phase may help trigger PTI, reflecting the plant’s attempt to resist the infection.

Furthermore, our results suggest that miR472-5p plays a complex role in regulating plant immunity depending on the phase of pathogen interaction. It is upregulated in NP vs. PA and BP vs. PA phases. Meanwhile, it is downregulated in the NP vs. BP phase. It has been observed that miR472a from poplar is involved in the defense against the necrotrophic fungus *Cystora chrysosperma* by targeting nucleotide-binding site (NBS) and leucine-rich (LRR) genes, which are negative regulators of necrotrophic resistance (Su et al. 2018). Hence, the increase in miR472-5p levels might be part of a regulatory response aimed at enhancing *A. thaliana*’s defense against the invading *C. higginsianum* during the initial stages of infection (PA and BP). The downregulation of miR472-5p in the NP phase compared to the BP phase could indicate a shift in the plant’s response to the pathogen as it transitions to a necrotrophic lifestyle. This suggests that *A. thaliana* may be less reliant on miR472-5p to manage NBS-LRR gene expression during this phase, which could mean that its defense strategy shifts at this point, possibly relying on other mechanisms to combat *C. higginsianum*.

miR165-5p was downregulated in the NP vs. BP but upregulated in NP vs. PA and BP vs. PA. miR165/166 have a role in auxin and abscisic acid (ABA) signaling, which are key players in stress responses in *Arabidopsis* (Jia et al. 2015). Hence, miR165-5p could be actively modulating *A. thaliana*’s immunity by regulating HD-ZIP III transcription factors across infection stages. During the early infection stages (PA/BP), its upregulation may suppress HD-ZIP III genes, inhibiting ABA-mediated defenses (Zhou et al. 2007), which creates a favorable environment for pathogen establishment as the plant is not yet defending itself. Conversely, its downregulation in NP could activate HD-ZIP III genes transcription, triggering ABA-responsive genes (like RD26) and auxin signaling to fight fungal spread (Zhou et al. 2007).

miR169 family members are known to play a role in responses to biotic/abiotic stress in poplar by targeting nuclear transcriptional factors Y subunit alfa (NF-YA) (Wang et al. 2022). The observed expression patterns of miR169-3p from our results demonstrate downregulation in BP vs. PA and NP vs. BP but an upregulation in NP vs. PA. Its upregulation in the NP vs. PA phase could mirror poplar’s stress response.

We did not detect differential accumulation of miRNA160, which has been reported to be upregulated in *Arabidopsis* infected with *P. syringe* and in cassava infected with *C. higginsianum* (Zhang et al. 2021; Pinweha et al. 2015). Other miRNA families, such as miR773, that have been previously identified to play a role in the plant defense mechanism of *Arabidopsis* against *C. higginsianum* were absent from our results (Salvador-Guirao et al. 2018). This lack of detection may be due to the absence of an uninfected *A. thaliana* control sample in our dataset. As a result, we could only assess differences across infection phases, but not between infected and uninfected states of the plant, where differential accumulation of these miRNAs might have been detected.

To better illustrate these host and pathogen dynamics, we summarized the phase-specific regulation of miRNAs and tRNA fragments in a schematic overview (Figure 4). This figure highlights how certain sRNAs are consistently regulated across all phases, while others display stage-specific patterns.

**Figure 4.**
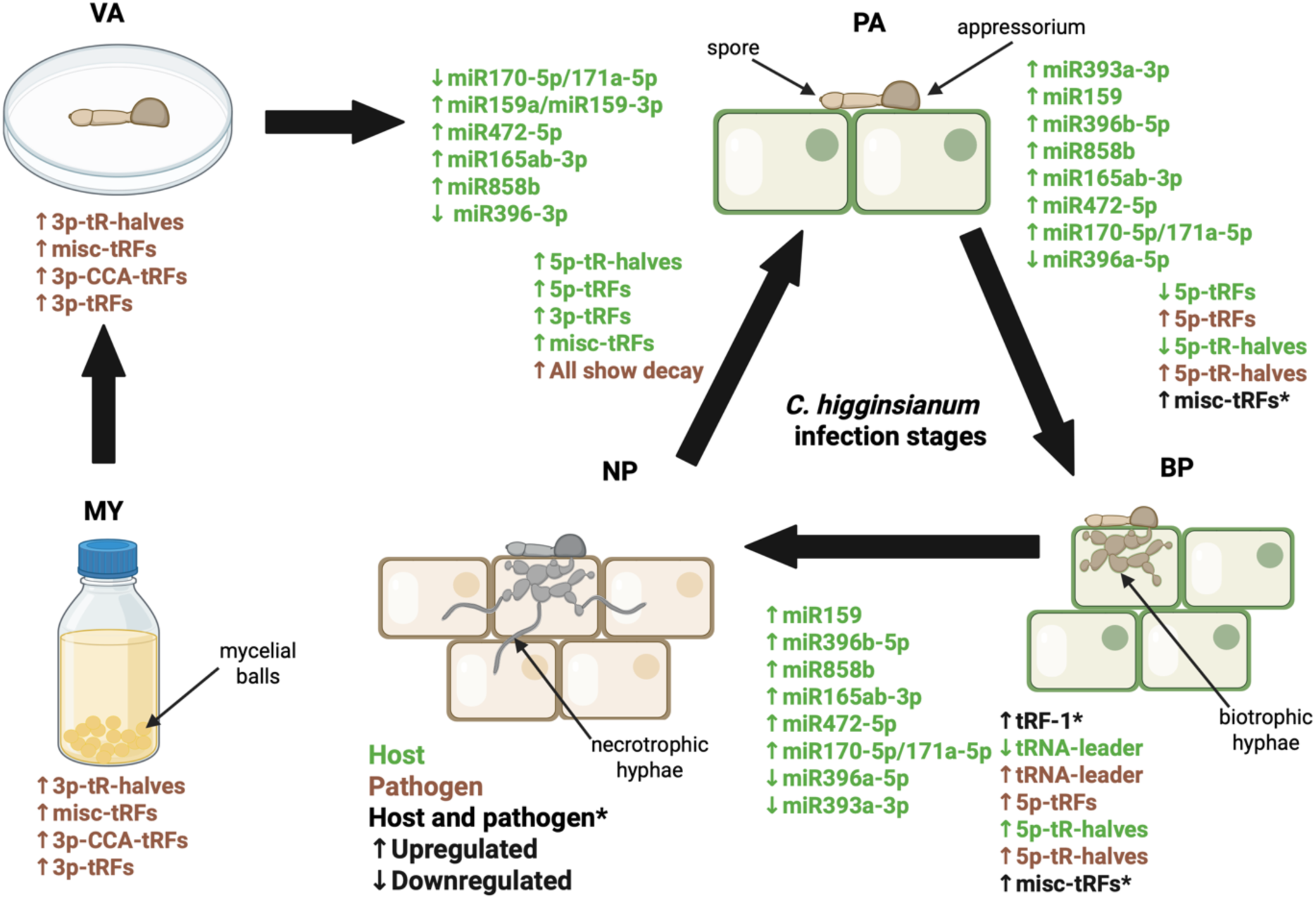
Schematic summary of miRNA and tRNA fragment dynamics during the infection stages of *C. higginsianum* in *A. thaliana*. The stages shown are: *In Planta* Appressorium (PA), Biotrophic Phase (BP), Necrotrophic Phase (NP), *In Vitro* Appressorium (VA), and Mycelia (MY). Differential accumulation of the sRNAs is color coded: green for *A. thaliana*, brown for *C. higginsianum*, and black for miRNAs and tRNA fragments that are similarly differential accumulation in both host and pathogen. Upregulation is indicated by upward arrows, and downregulation by downward arrows.

### tRFs as key players in infection dynamics

Our results show that for *C. higginsianum* 3p-tRNA halves are the most abundant tRFs across all infection phases, which could imply that they not only play an important role in fungal infection. Nevertheless, their abundance reduces considerably in the NP stage. In fact, in *C. higginsianum* all tRFs drop in abundance in the NP phase, suggesting RNA decay. 3p-CCA-tRFs have low abundance in most phases but peak in the MY phase, possibly linked to a specific fungal developmental stage. misc-tRFs, 5p-tRNA halves, and 5p-tRFs show higher abundance in early phases (PA, BP) indicating potential roles in early infection establishment. tRF-1 and tRNA leader peak in the BP phase, implying involvement in BP phase transitions, possibly contributing to its pathogenicity.

For *A. thaliana*, 3p-tRNA halves are consistently more abundant than in *C. higginsianum*, but both show a decreasing trend from PA to NP, suggesting a conserved response to infection. misc-tRFs, 5p-tRFs and 3p-tRFs increase exhibit opposite trends, it decreases in *C. higginsianum* but increases in *A. thaliana* (from PA to NP). This may reflect host vs. pathogen regulatory differences. 5p-tRNA halves peak in the NP phase in *A. thaliana* but remain lower in *C. higginsianum*, suggesting host-specific stress responses. 3p-CCA-tRFs, tRF-1 and tRNA leader are consistently in low in the *A. thaliana*, suggesting they might have minimal functional relevance to defense against the pathogen.

The contrasting trends (in misc-tRFs, 5p-tRFs and 3p-tRFs) suggest that *C. higginsianum* may downregulate certain tRFs in later infection stages, while *A. thaliana* upregulates them as a defense response. The high abundance of 3p-tRNA halves in both species implies a conserved role in RNA regulation during stress/infection (Chen et al. 2025; Pawar et al. 2020). The MY-phase peak of 3p-CCA-tRFs in *C. higginsianum* may correlate with fungal sporulation or invasive growth. However, we understand that ligation-based sRNA sequencing is known to have an intrinsic bias in capturing different tRNA fragments (Shi et al. 2021), and further studies of tRNA processing during plant-pathogen interactions are needed to validate this hypothesis.

In conclusion, the interaction between *C. higginsianum* and *A. thaliana* reflects a constant molecular tug-of-war mediated by small RNAs. In the early stages, the fungus deploys abundant 29 nt sRNAs, while the host counters with diverse 21–24 nt and 30 nt sRNAs, alongside stage-specific miRNA regulation (e.g., downregulating miR396 early, suppressing miR858b later). During the necrotrophic phase, fungal sRNAs decline sharply, likely reflecting enhanced RNA turnover and degradation associated with host cell death. At the same time, *Arabidopsis* shifts toward a dominant 21 nt sRNA population, but its overall regulatory capacity weakens as infection progresses. These results demonstrate that sRNA dynamics are not only central to the host–pathogen arms race but also highlight RNA turnover as a key mechanism shaping the outcome of infection.

## Acknowledgments

We would like to express our sincere appreciation to Dr. Corbin D. Jones for generously dedicating resources to support the successful completion of this work. This work was funded by <u>NSF grant IOS-2243536</u> to C.E.A. R.J.O. and T.-T.-H.C. were supported by funding from the Agence Nationale de la Recherche (ERA- CAPS grant ANR-17-CAPS-0004-01). The BIOGER unit benefits from the support of Saclay Plant Sciences (ANR-17-EUR-0007). This work was supported by National Science Foundation (NSF) grant IOS-2243536 to PB.

## Author Contributions

P.B., B. C. M. and R.J.O. designed the experiment. T.-T.-H.C. generated the plant-fungus samples and RNA isolation. P.B. generated the small RNA libraries. C.E.A analyzed data. P.B and C.E.A wrote the first draft of the manuscript, which all authors read, commented on, and edited.

## Data Availability Statement

Raw sRNA sequencing data from this article can be found in the NCBI Gene Expression Omnibus under accession number GSE159900.

## Supporting information

**Table S1.** Genomic coordinates of predicted rRNA genes from *C. higginsianum* genome.

**Table S2.** Predicted tRNA genes, tRNA pseudogenes, and atypical tRNA candidates from *C. higginsianum* genome.

**Table S3.** Classification and genomic distribution of repetitive elements from *C. higginsianum* genome.

**Figure S1.** Percent alignment (y-axis) bar plots of sRNA reads from *C. higginsianum* and *A. thaliana* across various infection phases. The phases represented for *C. higginsianum* growth include: *In Planta* Appressorium (PA), Biotrophic Phase (BP), Necrotrophic Phase (NP), *In Vitro* Appressorium (VA), and Mycelia (MY). Each phase encompasses three biological replicates (1, 2, and 3), indicated on the x-axis. Bars are color-coded by species, with *C. higginsianum* depicted in mustard yellow and *A. thaliana* in blue.

**Figure S2.** A) Abundance of sRNA reads from *C. higginsianum* across different infection phases based on size distribution. In the *x-axis* are the nucleotide size (pb) of the sRNA reads during the pathogen infection stages: *In Planta* Appressorium (PA), Biotrophic Phase (BP), Necrotrophic Phase (NP), *In Vitro* Appressorium (VA), and Mycelia (MY). Each phase includes three replicates: 1 (blue), 2 (mustard-yellow), 3 (orange). In the *y-axis*, abundance in RPM is depicted. B) Abundance of sRNA reads from *A. thaliana* infection from *C. higginsianum* across different infection phases of the pathogen (PA, BP and NP) based on size distribution. Each phase includes three replicates: 1 (blue), 2 (green), 3 (mustard-yellow).

**Figure S3.** Size distribution profiles of sRNAs in bp (*x-axis*) that mapped to rRNAs in *C. higginsianum*. The *y-axis* displays RNA abundance in reads per million (RPM) for the different stages of *C. higginsianum*: *In Planta* Appressorium (PA), Biotrophic Phase (BP), Necrotrophic Phase (NP), *In Vitro* Appressorium (VA), and Mycelia (MY). Three replicates are indicated by distinct colors (blue, mustard-yellow, and orange).

**Figure S4.** Size distribution profiles of sRNAs in bp (*x-axis*) that mapped to TEs in *C. higginsianum*. The *y-axis* displays RNA abundance in reads per million (RPM) for the different stages of *C. higginsianum*: *In Planta* Appressorium (PA), Biotrophic Phase (BP), Necrotrophic Phase (NP), *In Vitro* Appressorium (VA), and Mycelia (MY). Three replicates are indicated by distinct colors (blue, mustard-yellow, and orange).

**Figure S5.** Size distribution profiles of sRNAs in bp (*x-axis*) that mapped to genes in *C. higginsianum*. The *y-axis* displays RNA abundance in reads per million (RPM) for the different stages of *C. higginsianum*: *In Planta* Appressorium (PA), Biotrophic Phase (BP), Necrotrophic Phase (NP), *In Vitro* Appressorium (VA), and Mycelia (MY). Three replicates are indicated by distinct colors (blue, mustard-yellow, and orange).

**Figure S6.** Size distribution profiles of sRNAs in bp (*x-axis*) that mapped to tRNAs in *C. higginsianum*. The *y-axis* displays RNA abundance in reads per million (RPM) for the different stages of *C. higginsianum*: *In Planta* Appressorium (PA), Biotrophic Phase (BP), Necrotrophic Phase (NP), *In Vitro* Appressorium (VA), and Mycelia (MY). Three replicates are indicated by distinct colors (blue, mustard-yellow, and orange).

**Figure S7.** Size distribution profiles of sRNAs in bp (*x-axis*) that mapped to genes in *A. thaliana*. The *y-axis* displays RNA abundance in reads per million (RPM) for the different stages of *C. higginsianum*: *In Planta* Appressorium (PA), Biotrophic Phase (BP) and Necrotrophic Phase (NP). Three replicates are indicated by distinct colors (blue, green, and mustard-yellow).

**Figure S8.** Size distribution profiles of sRNAs in bp (*x-axis*) that mapped to rRNAs in *A. thaliana*. The *y-axis* displays RNA abundance in reads per million (RPM) for the different stages of *C. higginsianum*: *In Planta* Appressorium (PA), Biotrophic Phase (BP) and Necrotrophic Phase (NP). Three replicates are indicated by distinct colors (blue, green, and mustard-yellow).

**Figure S9.** Size distribution profiles of sRNAs in bp (*x-axis*) that mapped to ncRNAs in *A. thaliana*. The *y-axis* displays RNA abundance in reads per million (RPM) for the different stages of *C. higginsianum*: *In Planta* Appressorium (PA), Biotrophic Phase (BP) and Necrotrophic Phase (NP). Three replicates are indicated by distinct colors (blue, green, and mustard-yellow).

**Figure S10.** Size distribution profiles of sRNAs in bp (*x-axis*) that mapped to TEs in *A. thaliana*. The *y-axis* displays RNA abundance in reads per million (RPM) for the different stages of *C. higginsianum*: *In Planta* Appressorium (PA), Biotrophic Phase (BP) and Necrotrophic Phase (NP). Three replicates are indicated by distinct colors (blue, green, and mustard-yellow).

**Figure S11.** Size distribution profiles of sRNAs in bp (*x-axis*) that mapped to miRNAs in *A. thaliana*. The *y-axis* displays RNA abundance in reads per million (RPM) for the different stages of *C. higginsianum*: *In Planta* Appressorium (PA), Biotrophic Phase (BP) and Necrotrophic Phase (NP). Three replicates are indicated by distinct colors (blue, green, and mustard-yellow).

**Figure S12.** Size distribution profiles of sRNAs in bp (*x-axis*) that mapped to snRNAs in *A. thaliana*. The *y-axis* displays RNA abundance in reads per million (RPM) for the different stages of *C. higginsianum*: *In Planta* Appressorium (PA), Biotrophic Phase (BP) and Necrotrophic Phase (NP). Three replicates are indicated by distinct colors (blue, green, and mustard-yellow).

**Figure S13.** Size distribution profiles of sRNAs in bp (*x-axis*) that mapped to tRNAs in *A. thaliana*. The *y-axis* displays RNA abundance in reads per million (RPM) for the different stages of *C. higginsianum*: *In Planta* Appressorium (PA), Biotrophic Phase (BP) and Necrotrophic Phase (NP). Three replicates are indicated by distinct colors (blue, green, and mustard-yellow).

**Figure S14.** Size distribution profiles of sRNAs in bp (*x-axis*) that mapped to snoRNA in *A. thaliana*. The *y-axis* displays RNA abundance in reads per million (RPM) for the different stages of *C. higginsianum*: *In Planta* Appressorium (PA), Biotrophic Phase (BP) and Necrotrophic Phase (NP). Three replicates are indicated by distinct colors (blue, green, and mustard-yellow).

