## Supplemental Figures and Tables for "Stage-Specific RNA Turnover Drives Small RNA Dynamics in *Arabidopsis* – *Colletotrichum* Interactions"

### Supplementary Material

**Table S1.** Genomic coordinates of predicted rRNA genes from *C. higginsianum* genome.

| Sequence name | Source | Feature | Start | End | Score | +/- | Frame | Attribute |
|---|---|---|---|---|---|---|---|---|
| unitig_22 | RNAmmer-1.2 | rRNA | 10958 | 14482 | 3223.6 | - | . | 28s_rRNA |
| unitig_17 | RNAmmer-1.2 | rRNA | 15792 | 19317 | 3239.7 | - | . | 28s_rRNA |
| unitig_18 | RNAmmer-1.2 | rRNA | 23140 | 26656 | 3203.8 | - | . | 28s_rRNA |
| unitig_25 | RNAmmer-1.2 | rRNA | 5358 | 8883 | 3239.7 | + | . | 28s_rRNA |
| unitig_17 | RNAmmer-1.2 | rRNA | 7628 | 11153 | 3239.7 | - | . | 28s_rRNA |
| unitig_20 | RNAmmer-1.2 | rRNA | 7704 | 11229 | 3239.7 | + | . | 28s_rRNA |
| unitig_17 | RNAmmer-1.2 | rRNA | 23953 | 27475 | 3082.7 | - | . | 28s_rRNA |
| unitig_15 | RNAmmer-1.2 | rRNA | 12385 | 15910 | 3239.7 | - | . | 28s_rRNA |
| unitig_19 | RNAmmer-1.2 | rRNA | 19568 | 23093 | 3239.7 | - | . | 28s_rRNA |
| chromosome_7 | RNAmmer-1.2 | rRNA | 52 | 3576 | 3205 | - | . | 28s_rRNA |
| unitig_15 | RNAmmer-1.2 | rRNA | 4240 | 7765 | 3239.7 | - | . | 28s_rRNA |
| unitig_19 | RNAmmer-1.2 | rRNA | 3163 | 6688 | 3239.7 | - | . | 28s_rRNA |
| unitig_14 | RNAmmer-1.2 | rRNA | 21984 | 25509 | 3239.7 | - | . | 28s_rRNA |
| unitig_18 | RNAmmer-1.2 | rRNA | 6712 | 10237 | 3239.7 | - | . | 28s_rRNA |
| unitig_16 | RNAmmer-1.2 | rRNA | 18122 | 21647 | 3239.7 | + | . | 28s_rRNA |
| unitig_23 | RNAmmer-1.2 | rRNA | 4945 | 8470 | 3239.7 | + | . | 28s_rRNA |
| unitig_16 | RNAmmer-1.2 | rRNA | 9931 | 13456 | 3239.7 | + | . | 28s_rRNA |
| unitig_19 | RNAmmer-1.2 | rRNA | 11354 | 14879 | 3239.7 | - | . | 28s_rRNA |
| unitig_24 | RNAmmer-1.2 | rRNA | 4596 | 8115 | 3182.4 | - | . | 28s_rRNA |
| unitig_18 | RNAmmer-1.2 | rRNA | 14926 | 18451 | 3239.7 | - | . | 28s_rRNA |
| unitig_22 | RNAmmer-1.2 | rRNA | 2782 | 6304 | 3231.2 | - | . | 28s_rRNA |
| unitig_21 | RNAmmer-1.2 | rRNA | 15947 | 19472 | 3239.7 | + | . | 28s_rRNA |
| unitig_24 | RNAmmer-1.2 | rRNA | 12743 | 16244 | 2990.2 | - | . | 28s_rRNA |
| chromosome_7 | RNAmmer-1.2 | rRNA | 8240 | 11765 | 3239.7 | - | . | 28s_rRNA |
| unitig_21 | RNAmmer-1.2 | rRNA | 7733 | 11258 | 3239.7 | + | . | 28s_rRNA |
| unitig_16 | RNAmmer-1.2 | rRNA | 1763 | 5288 | 3239.7 | + | . | 28s_rRNA |
| unitig_14 | RNAmmer-1.2 | rRNA | 30175 | 33701 | 3241.3 | - | . | 28s_rRNA |
| unitig_28 | RNAmmer-1.2 | rRNA | 11463 | 14981 | 3205.3 | - | . | 28s_rRNA |
| unitig_22 | RNAmmer-1.2 | rRNA | 19135 | 22645 | 3156 | - | . | 28s_rRNA |
| unitig_20 | RNAmmer-1.2 | rRNA | 15895 | 19420 | 3239.7 | + | . | 28s_rRNA |
| unitig_28 | RNAmmer-1.2 | rRNA | 3282 | 6805 | 3230.5 | - | . | 28s_rRNA |
| unitig_17 | RNAmmer-1.2 | rRNA | 1 | 2964 | 1900.6 | - | . | 28s_rRNA |
| unitig_20 | RNAmmer-1.2 | rRNA | 2 | 3038 | 2743.8 | + | . | 28s_rRNA |
| unitig_21 | RNAmmer-1.2 | rRNA | 3 | 3044 | 2755.7 | + | . | 28s_rRNA |
| unitig_25 | RNAmmer-1.2 | rRNA | 13526 | 17051 | 3239.7 | + | . | 28s_rRNA |
| unitig_23 | RNAmmer-1.2 | rRNA | 13056 | 16581 | 3239.7 | + | . | 28s_rRNA |

| | | | | | | | | |
|---|---|---|---|---|---|---|---|---|
| chromosome_7 | RNAmmer-1.2 | rRNA | 16454 | 19979 | 3239.7 | - | . | 28s_rRNA |
| unitig_15 | RNAmmer-1.2 | rRNA | 20594 | 24119 | 3239.7 | - | . | 28s_rRNA |
| unitig_28 | RNAmmer-1.2 | rRNA | 15163 | 16992 | 1414.7 | - | . | 18s_rRNA |
| unitig_16 | RNAmmer-1.2 | rRNA | 7951 | 9745 | 1439.1 | + | . | 18s_rRNA |
| unitig_20 | RNAmmer-1.2 | rRNA | 5724 | 7518 | 1439.1 | + | . | 18s_rRNA |
| unitig_14 | RNAmmer-1.2 | rRNA | 25695 | 27489 | 1439.1 | - | . | 18s_rRNA |
| unitig_22 | RNAmmer-1.2 | rRNA | 14668 | 16460 | 1429.7 | - | . | 18s_rRNA |
| unitig_16 | RNAmmer-1.2 | rRNA | 3 | 1577 | 1184.4 | + | . | 18s_rRNA |
| unitig_18 | RNAmmer-1.2 | rRNA | 2223 | 4017 | 1439.1 | - | . | 18s_rRNA |
| unitig_20 | RNAmmer-1.2 | rRNA | 22104 | 23898 | 1439.1 | + | . | 18s_rRNA |
| chromosome_7 | RNAmmer-1.2 | rRNA | 11951 | 13745 | 1439.1 | - | . | 18s_rRNA |
| unitig_23 | RNAmmer-1.2 | rRNA | 11076 | 12870 | 1439.1 | + | . | 18s_rRNA |
| unitig_19 | RNAmmer-1.2 | rRNA | 15065 | 16859 | 1439.1 | - | . | 18s_rRNA |
| unitig_25 | RNAmmer-1.2 | rRNA | 3378 | 5172 | 1439.1 | + | . | 18s_rRNA |
| unitig_24 | RNAmmer-1.2 | rRNA | 8300 | 10078 | 1298.5 | - | . | 18s_rRNA |
| unitig_21 | RNAmmer-1.2 | rRNA | 5753 | 7547 | 1439.1 | + | . | 18s_rRNA |
| unitig_16 | RNAmmer-1.2 | rRNA | 24333 | 26127 | 1439.1 | + | . | 18s_rRNA |
| chromosome_7 | RNAmmer-1.2 | rRNA | 20165 | 21959 | 1439.1 | - | . | 18s_rRNA |
| unitig_15 | RNAmmer-1.2 | rRNA | 7951 | 9745 | 1439.1 | - | . | 18s_rRNA |
| unitig_25 | RNAmmer-1.2 | rRNA | 11546 | 13340 | 1439.1 | + | . | 18s_rRNA |
| unitig_28 | RNAmmer-1.2 | rRNA | 6990 | 8780 | 1406.2 | - | . | 18s_rRNA |
| unitig_18 | RNAmmer-1.2 | rRNA | 10423 | 12217 | 1439.1 | - | . | 18s_rRNA |
| unitig_21 | RNAmmer-1.2 | rRNA | 13967 | 15761 | 1439.1 | + | . | 18s_rRNA |
| unitig_17 | RNAmmer-1.2 | rRNA | 11339 | 13133 | 1439.1 | - | . | 18s_rRNA |
| unitig_23 | RNAmmer-1.2 | rRNA | 19290 | 21084 | 1439.1 | + | . | 18s_rRNA |
| unitig_15 | RNAmmer-1.2 | rRNA | 24305 | 26099 | 1439.1 | - | . | 18s_rRNA |
| unitig_23 | RNAmmer-1.2 | rRNA | 2965 | 4759 | 1439.1 | + | . | 18s_rRNA |
| unitig_14 | RNAmmer-1.2 | rRNA | 16850 | 19322 | 127.2 | - | . | 18s_rRNA |
| unitig_19 | RNAmmer-1.2 | rRNA | 23279 | 25073 | 1439.1 | - | . | 18s_rRNA |
| unitig_17 | RNAmmer-1.2 | rRNA | 3148 | 4942 | 1439.1 | - | . | 18s_rRNA |
| unitig_15 | RNAmmer-1.2 | rRNA | 16095 | 17889 | 1439.1 | - | . | 18s_rRNA |
| unitig_15 | RNAmmer-1.2 | rRNA | 9 | 1555 | 999.9 | - | . | 18s_rRNA |
| unitig_22 | RNAmmer-1.2 | rRNA | 6489 | 8282 | 1439.2 | - | . | 18s_rRNA |
| unitig_20 | RNAmmer-1.2 | rRNA | 13915 | 15709 | 1439.1 | + | . | 18s_rRNA |
| chromosome_7 | RNAmmer-1.2 | rRNA | 3761 | 5555 | 1439.1 | - | . | 18s_rRNA |
| unitig_19 | RNAmmer-1.2 | rRNA | 6874 | 8668 | 1439.1 | - | . | 18s_rRNA |
| unitig_17 | RNAmmer-1.2 | rRNA | 19502 | 21295 | 1432 | - | . | 18s_rRNA |
| unitig_16 | RNAmmer-1.2 | rRNA | 16142 | 17936 | 1439.1 | + | . | 18s_rRNA |
| unitig_24 | RNAmmer-1.2 | rRNA | 16426 | 18210 | 1358.7 | - | . | 18s_rRNA |
| unitig_21 | RNAmmer-1.2 | rRNA | 22112 | 23906 | 1439.1 | + | . | 18s_rRNA |

| | | | | | | | | |
|---|---|---|---|---|---|---|---|---|
| unitig_18 | RNAmmer-1.2 | rRNA | 18637 | 20431 | 1439.1 | - | . | 18s_rRNA |
| unitig_24 | RNAmmer-1.2 | rRNA | 146 | 1928 | 1316.1 | - | . | 18s_rRNA |
| chromosome_7 | RNAmmer-1.2 | rRNA | 1024226 | 1024341 | 54.1 | + | . | 8s_rRNA |
| chromosome_3 | RNAmmer-1.2 | rRNA | 3073219 | 3073334 | 55.5 | + | . | 8s_rRNA |
| chromosome_9 | RNAmmer-1.2 | rRNA | 2111753 | 2111868 | 53.6 | - | . | 8s_rRNA |
| chromosome_4 | RNAmmer-1.2 | rRNA | 3852044 | 3852159 | 59.1 | + | . | 8s_rRNA |
| chromosome_8 | RNAmmer-1.2 | rRNA | 3574044 | 3574159 | 58.3 | + | . | 8s_rRNA |
| chromosome_2 | RNAmmer-1.2 | rRNA | 2951514 | 2951629 | 56.7 | - | . | 8s_rRNA |
| chromosome_1 | RNAmmer-1.2 | rRNA | 4718404 | 4718519 | 55.6 | - | . | 8s_rRNA |
| chromosome_5 | RNAmmer-1.2 | rRNA | 2543028 | 2543143 | 58.3 | + | . | 8s_rRNA |
| chromosome_5 | RNAmmer-1.2 | rRNA | 2704868 | 2704983 | 53.6 | + | . | 8s_rRNA |
| chromosome_5 | RNAmmer-1.2 | rRNA | 3308236 | 3308351 | 58.3 | - | . | 8s_rRNA |
| chromosome_4 | RNAmmer-1.2 | rRNA | 3017612 | 3017727 | 59.1 | + | . | 8s_rRNA |
| chromosome_4 | RNAmmer-1.2 | rRNA | 2951811 | 2951926 | 54.1 | - | . | 8s_rRNA |
| chromosome_9 | RNAmmer-1.2 | rRNA | 636245 | 636360 | 55.7 | - | . | 8s_rRNA |
| chromosome_9 | RNAmmer-1.2 | rRNA | 2735137 | 2735252 | 56.4 | + | . | 8s_rRNA |
| chromosome_9 | RNAmmer-1.2 | rRNA | 403537 | 403652 | 57.2 | + | . | 8s_rRNA |
| chromosome_1 | RNAmmer-1.2 | rRNA | 3268296 | 3268411 | 46.7 | - | . | 8s_rRNA |
| chromosome_7 | RNAmmer-1.2 | rRNA | 3841105 | 3841220 | 53.5 | - | . | 8s_rRNA |
| chromosome_6 | RNAmmer-1.2 | rRNA | 2967022 | 2967137 | 58.3 | + | . | 8s_rRNA |
| chromosome_3 | RNAmmer-1.2 | rRNA | 1060937 | 1061052 | 54.5 | + | . | 8s_rRNA |
| chromosome_2 | RNAmmer-1.2 | rRNA | 1400369 | 1400484 | 50 | + | . | 8s_rRNA |
| chromosome_9 | RNAmmer-1.2 | rRNA | 2164642 | 2164757 | 55.6 | + | . | 8s_rRNA |
| chromosome_5 | RNAmmer-1.2 | rRNA | 3079589 | 3079704 | 37.6 | + | . | 8s_rRNA |
| chromosome_6 | RNAmmer-1.2 | rRNA | 495219 | 495334 | 41.6 | + | . | 8s_rRNA |
| chromosome_6 | RNAmmer-1.2 | rRNA | 2904069 | 2904184 | 57.2 | - | . | 8s_rRNA |
| chromosome_9 | RNAmmer-1.2 | rRNA | 3055263 | 3055378 | 58.3 | + | . | 8s_rRNA |
| chromosome_4 | RNAmmer-1.2 | rRNA | 4950725 | 4950840 | 39.4 | + | . | 8s_rRNA |
| chromosome_4 | RNAmmer-1.2 | rRNA | 4738912 | 4739027 | 55.6 | - | . | 8s_rRNA |
| chromosome_10 | RNAmmer-1.2 | rRNA | 2548472 | 2548587 | 50.5 | + | . | 8s_rRNA |
| chromosome_5 | RNAmmer-1.2 | rRNA | 2505802 | 2505917 | 53.6 | - | . | 8s_rRNA |
| chromosome_1 | RNAmmer-1.2 | rRNA | 3393062 | 3393177 | 52.1 | + | . | 8s_rRNA |
| chromosome_4 | RNAmmer-1.2 | rRNA | 2973039 | 2973154 | 55.6 | + | . | 8s_rRNA |
| chromosome_4 | RNAmmer-1.2 | rRNA | 3263165 | 3263280 | 55.6 | - | . | 8s_rRNA |
| chromosome_10 | RNAmmer-1.2 | rRNA | 2545666 | 2545781 | 52.1 | - | . | 8s_rRNA |
| chromosome_9 | RNAmmer-1.2 | rRNA | 1084373 | 1084488 | 58.3 | - | . | 8s_rRNA |
| chromosome_10 | RNAmmer-1.2 | rRNA | 1258520 | 1258635 | 55.6 | + | . | 8s_rRNA |
| chromosome_2 | RNAmmer-1.2 | rRNA | 1950064 | 1950179 | 60.8 | + | . | 8s_rRNA |
| chromosome_2 | RNAmmer-1.2 | rRNA | 2366925 | 2367040 | 61.8 | - | . | 8s_rRNA |
| chromosome_6 | RNAmmer-1.2 | rRNA | 3784946 | 3785061 | 53.6 | + | . | 8s_rRNA |

| | | | | | | | | |
|---|---|---|---|---|---|---|---|---|
| chromosome_6 | RNAmmer-1.2 | rRNA | 4008950 | 4009065 | 52.7 | + | . | 8s_rRNA |
| chromosome_12 | RNAmmer-1.2 | rRNA | 285030 | 285145 | 47.3 | - | . | 8s_rRNA |
| chromosome_1 | RNAmmer-1.2 | rRNA | 2568496 | 2568611 | 40.7 | + | . | 8s_rRNA |
| chromosome_7 | RNAmmer-1.2 | rRNA | 570895 | 571010 | 50.7 | - | . | 8s_rRNA |
| chromosome_1 | RNAmmer-1.2 | rRNA | 4430301 | 4430416 | 58.3 | - | . | 8s_rRNA |
| chromosome_4 | RNAmmer-1.2 | rRNA | 1452562 | 1452677 | 58.3 | - | . | 8s_rRNA |
| chromosome_9 | RNAmmer-1.2 | rRNA | 3261186 | 3261301 | 55.6 | - | . | 8s_rRNA |
| chromosome_7 | RNAmmer-1.2 | rRNA | 348296 | 348411 | 59.1 | + | . | 8s_rRNA |
| chromosome_2 | RNAmmer-1.2 | rRNA | 5523006 | 5523121 | 58.3 | - | . | 8s_rRNA |
| chromosome_8 | RNAmmer-1.2 | rRNA | 456204 | 456319 | 56.2 | - | . | 8s_rRNA |
| chromosome_12 | RNAmmer-1.2 | rRNA | 551677 | 551792 | 54.8 | - | . | 8s_rRNA |
| chromosome_2 | RNAmmer-1.2 | rRNA | 2510001 | 2510116 | 51.5 | + | . | 8s_rRNA |
| chromosome_5 | RNAmmer-1.2 | rRNA | 3412868 | 3412983 | 54.8 | - | . | 8s_rRNA |
| chromosome_2 | RNAmmer-1.2 | rRNA | 3104886 | 3105001 | 58.3 | + | . | 8s_rRNA |
| chromosome_2 | RNAmmer-1.2 | rRNA | 1334059 | 1334174 | 51.5 | - | . | 8s_rRNA |
| chromosome_9 | RNAmmer-1.2 | rRNA | 2787555 | 2787670 | 58.3 | - | . | 8s_rRNA |
| chromosome_10 | RNAmmer-1.2 | rRNA | 1483854 | 1483969 | 58.3 | + | . | 8s_rRNA |
| chromosome_9 | RNAmmer-1.2 | rRNA | 3291006 | 3291121 | 54.8 | + | . | 8s_rRNA |
| chromosome_4 | RNAmmer-1.2 | rRNA | 3214435 | 3214550 | 55.6 | - | . | 8s_rRNA |
| chromosome_6 | RNAmmer-1.2 | rRNA | 4052477 | 4052592 | 52.1 | - | . | 8s_rRNA |
| chromosome_9 | RNAmmer-1.2 | rRNA | 1791729 | 1791844 | 53.2 | + | . | 8s_rRNA |
| chromosome_12 | RNAmmer-1.2 | rRNA | 466911 | 467028 | 43.9 | + | . | 8s_rRNA |
| chromosome_8 | RNAmmer-1.2 | rRNA | 3891441 | 3891556 | 58.3 | + | . | 8s_rRNA |
| chromosome_7 | RNAmmer-1.2 | rRNA | 981311 | 981426 | 55.9 | + | . | 8s_rRNA |
| chromosome_10 | RNAmmer-1.2 | rRNA | 2989474 | 2989589 | 58.3 | + | . | 8s_rRNA |
| chromosome_10 | RNAmmer-1.2 | rRNA | 2552498 | 2552613 | 53.2 | - | . | 8s_rRNA |
| chromosome_5 | RNAmmer-1.2 | rRNA | 1724473 | 1724588 | 59.1 | - | . | 8s_rRNA |
| chromosome_1 | RNAmmer-1.2 | rRNA | 976137 | 976252 | 49.5 | + | . | 8s_rRNA |
| chromosome_10 | RNAmmer-1.2 | rRNA | 2195751 | 2195859 | 25.8 | + | . | 8s_rRNA |

**Table S2.** Predicted tRNA genes, tRNA pseudogenes, and atypical tRNA candidates from *C. higginsianum* genome.

| Category | Count |
|---|---|
| tRNAs decoding Standard 20 AA | 346 |
| Selenocysteine tRNAs (TCA) | 0 |
| Possible suppressor tRNAs (CTA,TTA,TCA) | 0 |
| tRNAs with undetermined/unknown isotypes | 1 |
| Predicted pseudogenes | 7 |
| **Total tRNAs** | 354 |

**Table S3.** Classification and genomic distribution of repetitive elements from *C. higginsianum* genome.

| | Category | Subcategory | Number of elements* | Length occupied (bp) | Percentage of sequence |
|---|---|---|---|---|---|
| Interspersed repeats | Retroelements | SINEs | 22 | 2015 | 0.00% |
| | | Penelope | 0 | 0 | 0.00% |
| | | LINEs | 28 | 88484 | 0.17% |
| | | CRE/SLACS | 0 | 0 | 0.00% |
| | | L2/CR1/Rex | 0 | 0 | 0.00% |
| | | R1/LOA/Jockey | 0 | 0 | 0.00% |
| | | R2/R4/NeSL | 0 | 0 | 0.00% |
| | | RTE/Bov-B | 0 | 0 | 0.00% |
| | | L1/CIN4 | 0 | 0 | 0.00% |
| | | LTR elements | 640 | 923177 | 1.82% |
| | | BEL/Pao | 0 | 0 | 0.00% |
| | | Ty1/Copia | 228 | 361268 | 0.71% |
| | | Gypsy/DIRS1 | 125 | 478313 | 0.94% |
| | | Retroviral | 0 | 0 | 0.00% |
| | | | **690** | **1013676** | **2.00%** |
| | DNA transposons | hobo-Activator | 0 | 0 | 0.00% |
| | | Tc1-IS630-Pogo | 892 | 1004920 | 1.98% |
| | | En-Spm | 0 | 0 | 0.00% |
| | | MULE-MuDR | 0 | 0 | 0.00% |
| | | PiggyBac | 0 | 0 | 0.00% |
| | | Tourist/Harbinger | 0 | 0 | 0.00% |
| | | Other (Mirage, P-element, Transib) | 0 | 0 | 0.00% |
| | | | **1069** | **1308038** | **2.58%** |
| | **Rolling-circles** | | **24** | **3251** | **0.01%** |
| | **Unclassified** | | **1910** | **485017** | **0.96%** |
| | | **Total** | | **2806731** | **5.53%** |
| Non interspersed repeats | **Small RNA** | | 182 | 349756 | 0.69% |
| | **Satellites** | | 0 | 0 | 0.00% |
| | **Simple repeats** | | 21475 | 822134 | 1.62% |
| | **Low complexity** | | 2212 | 104250 | 0.21% |
| | | **Total** | | **1276140** | **2.52%** |

*Most repeats fragmented by insertions or deletions have been counted as one element.

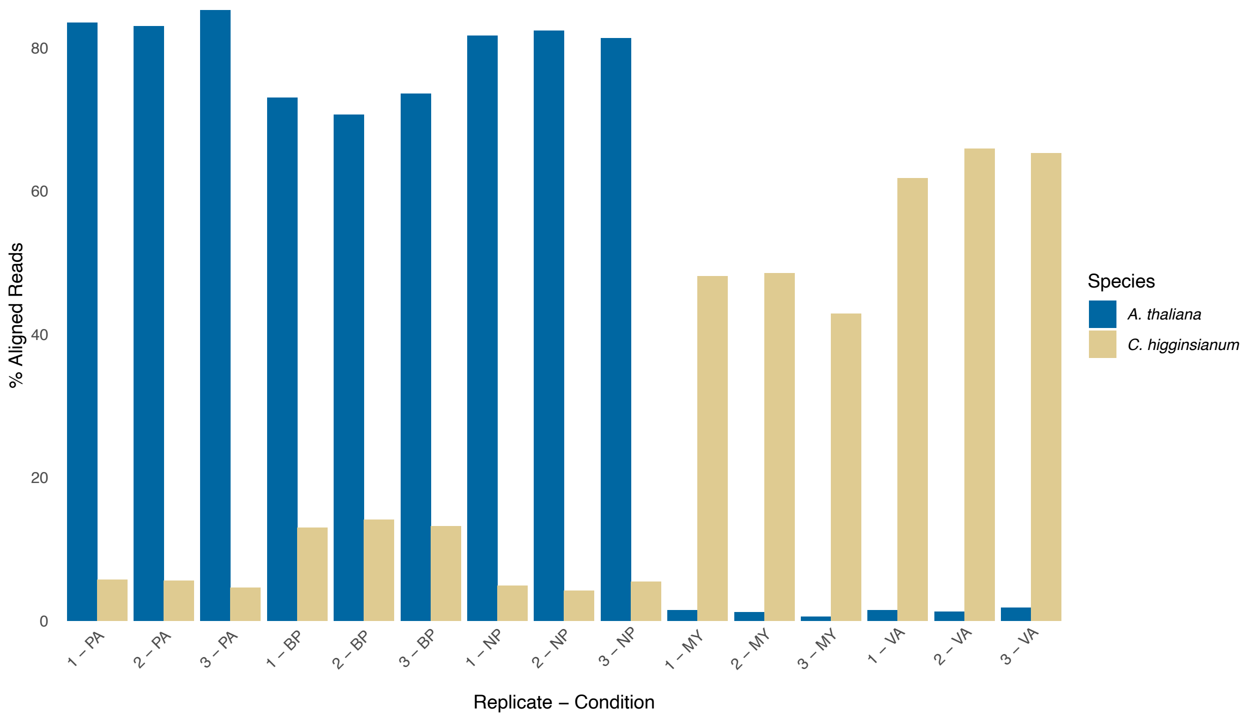


**Figure S1. Percent alignment (y-axis) bar plots of sRNA reads from *C. higginsianum* and *A. thaliana* across various infection stages.** The y-axis represents the percentage of aligned sRNA reads, while the x-axis shows the different stages of *C. higginsianum* development: In Planta Appressorium (PA), Biotrophic Phase (BP), Necrotrophic Phase (NP), In Vitro Appressorium (VA), and Mycelia (MY). Three biological replicates (1, 2, and 3) are displayed for each stage. Bars corresponding to *C. higginsianum* are shown in mustard yellow, and those corresponding to *A. thaliana* are shown in blue.

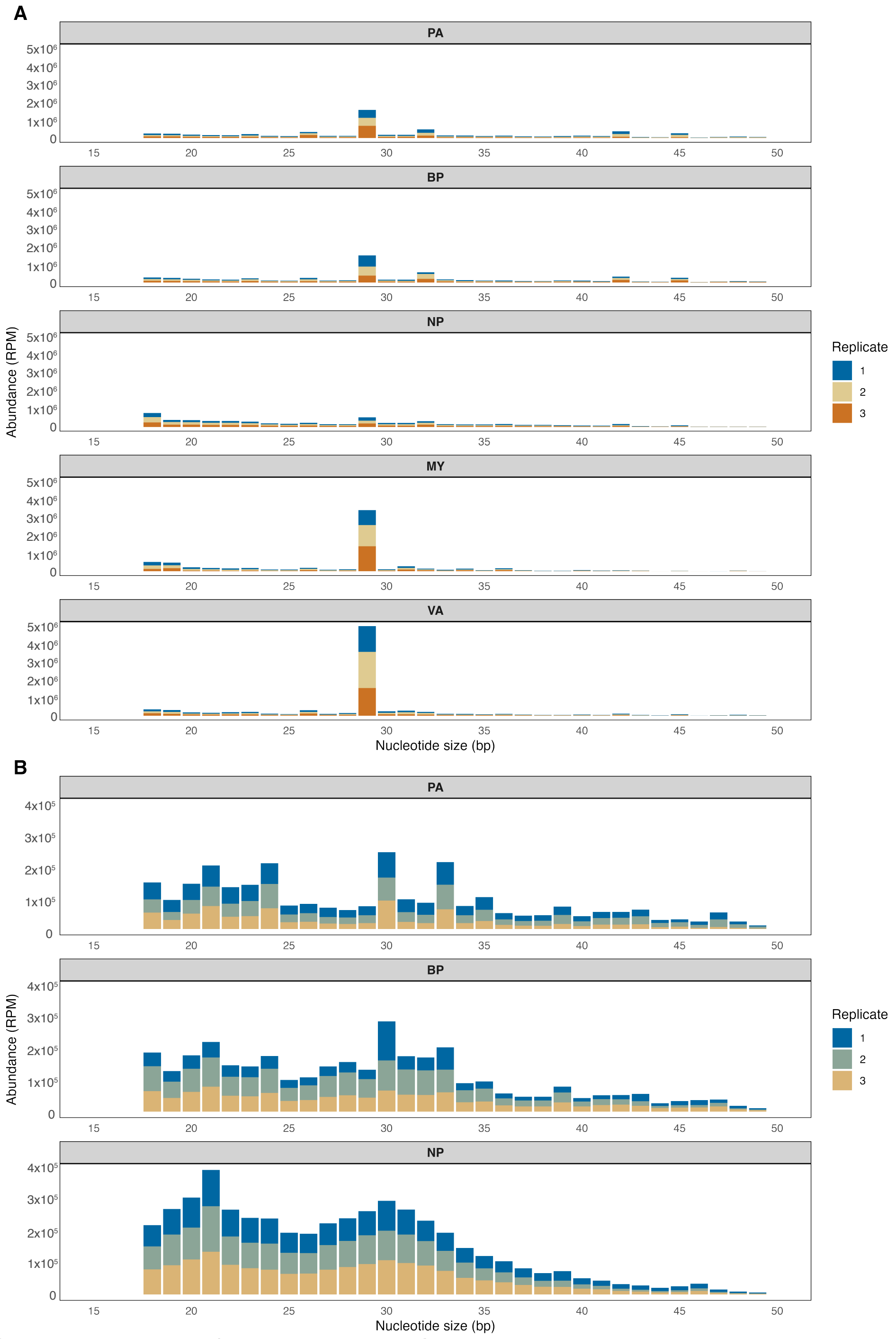


**Figure S2. Size distribution of sRNA reads in *C. higginsianum* and *A. thaliana* during infection.** A) Abundance of *C. higginsianum* sRNA reads across developmental and infection stages based on nucleotide length. The x-axis represents sRNA size (nt), and the y-axis shows read abundance in reads per million (RPM). Stages include In Planta Appressorium (PA), Biotrophic Phase (BP), Necrotrophic Phase (NP), In Vitro Appressorium (VA), and Mycelia (MY). Three biological replicates are represented by distinct colors: blue (1), mustard-yellow (2), and orange (3). B) Abundance of *A. thaliana* sRNA reads during infection by *C. higginsianum* at the PA, BP, and NP stages, shown as a function of nucleotide length (x-axis) and read abundance in RPM (y-axis). Three biological replicates are displayed in different colors: blue (1), green (2), and mustard-yellow (3).

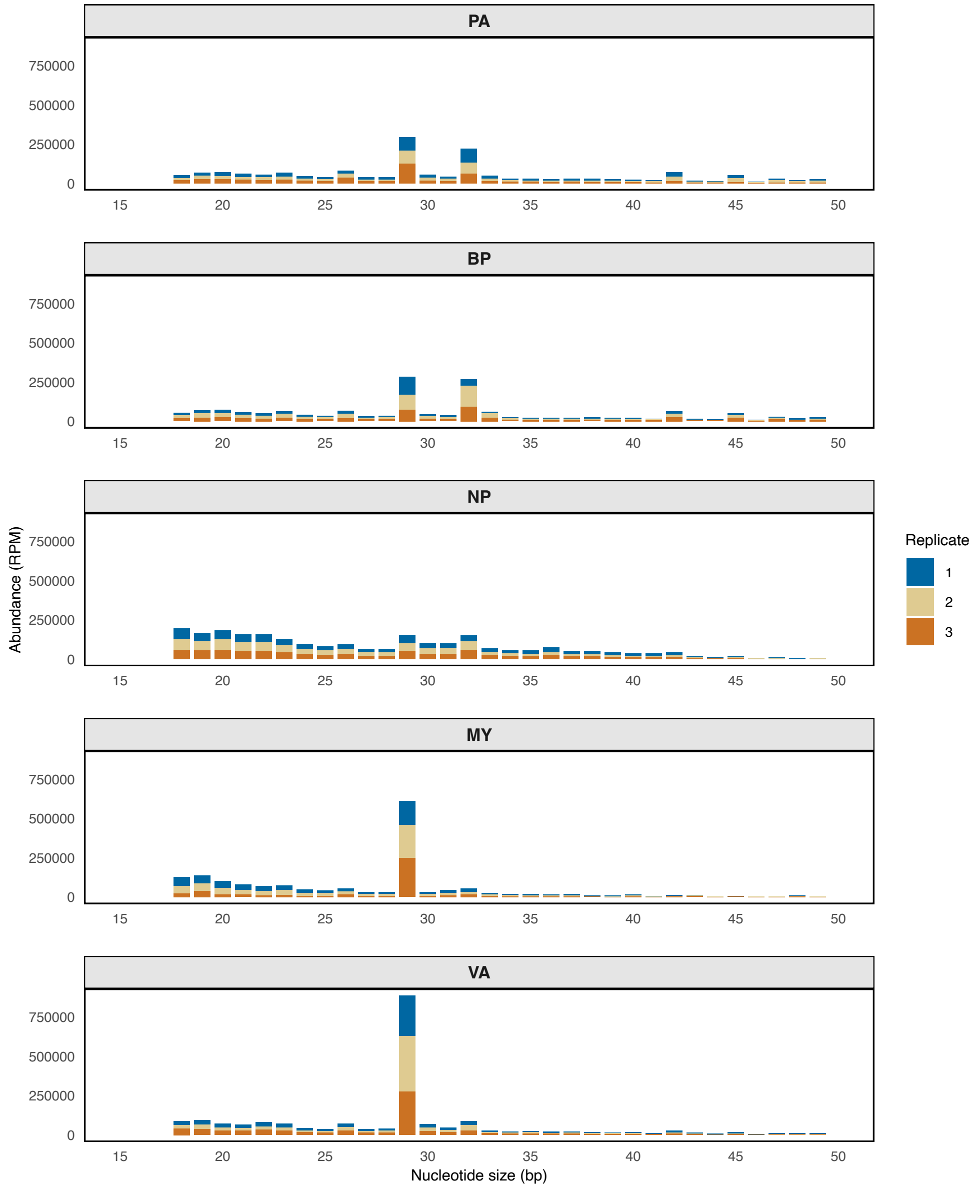


**Figure S3. Size distribution of rRNA-derived sRNAs in *C. higginsianum* across developmental and infection stages.**
The x-axis represents sRNA length (nt), and the y-axis shows RNA abundance in reads per million (RPM). Stages include In Planta Appressorium (PA), Biotrophic Phase (BP), Necrotrophic Phase (NP), In Vitro Appressorium (VA), and Mycelia (MY). Three biological replicates are represented by distinct colors: blue, mustard-yellow, and orange.

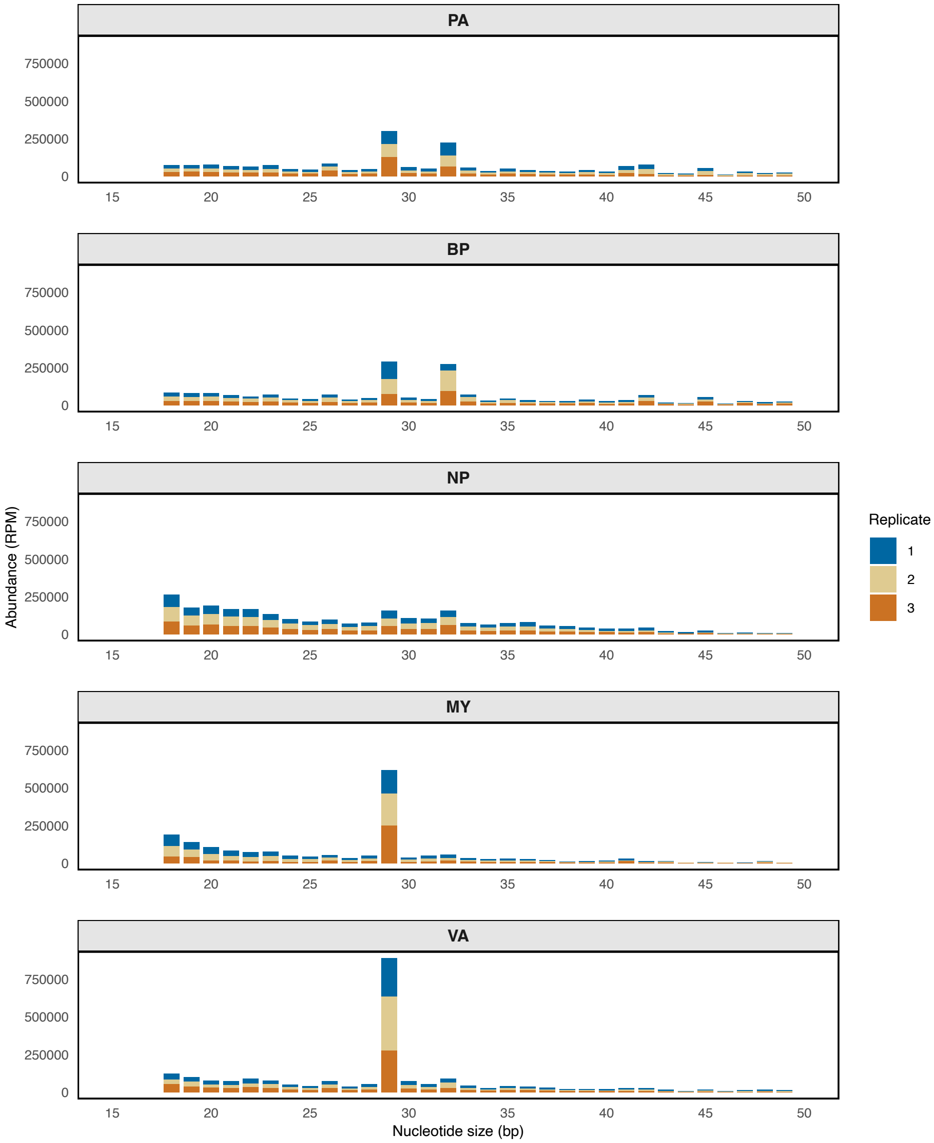


**Figure S4. Size distribution of TE-derived sRNAs in *C. higginsianum* across developmental and infection stages.**
The x-axis represents sRNA length (nt), and the y-axis shows RNA abundance in reads per million (RPM). Stages include In Planta Appressorium (PA), Biotrophic Phase (BP), Necrotrophic Phase (NP), In Vitro Appressorium (VA), and Mycelia (MY). Three biological replicates are represented by distinct colors: blue, mustard-yellow, and orange.

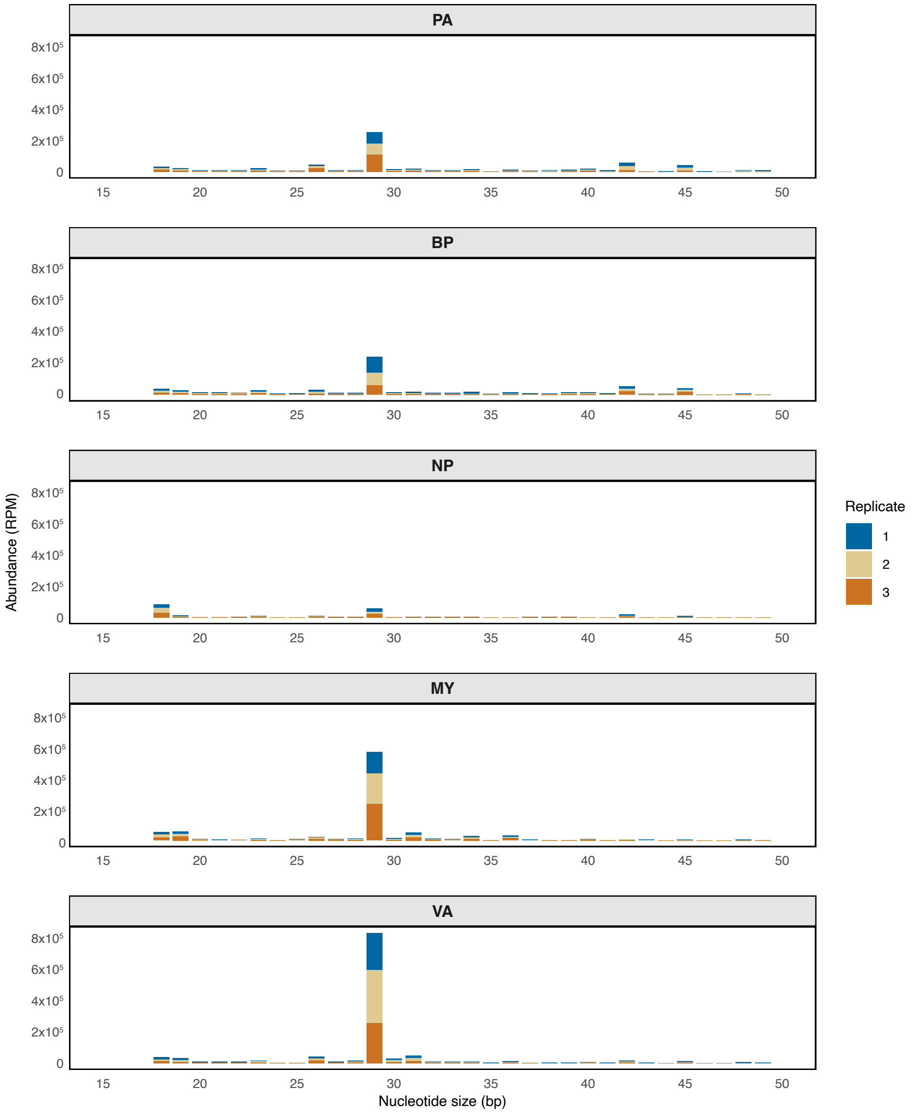


**Figure S5. Size distribution of gene-derived sRNAs in *C. higginsianum* across developmental and infection stages.** The x-axis represents sRNA length (nt), and the y-axis shows RNA abundance in reads per million (RPM). Stages include In Planta Appressorium (PA), Biotrophic Phase (BP), Necrotrophic Phase (NP), In Vitro Appressorium (VA), and Mycelia (MY). Three biological replicates are represented by distinct colors: blue, mustard-yellow, and orange.

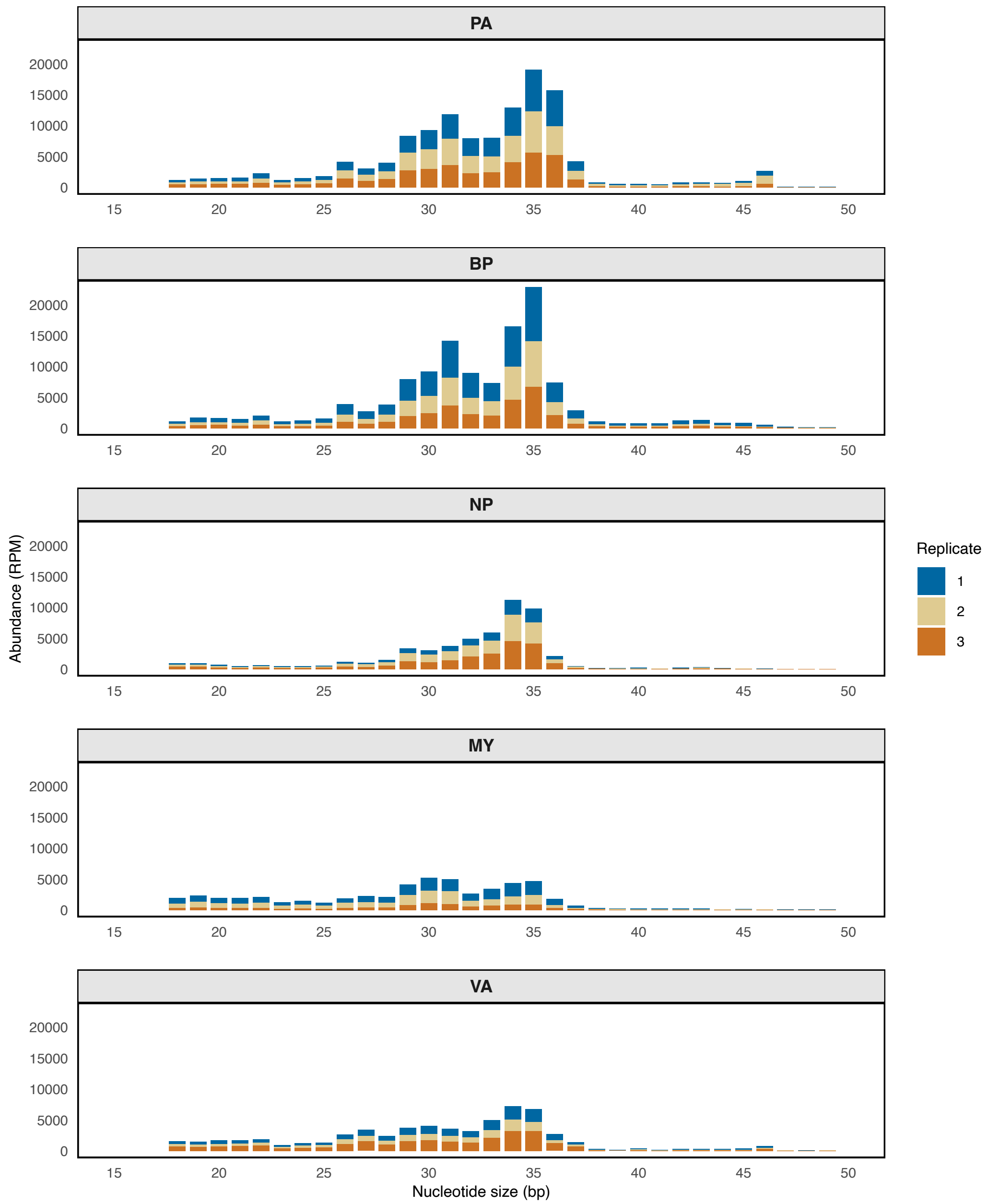


**Figure S6. Size distribution of tRNA-derived sRNAs in *C. higginsianum* across developmental and infection stages.**

The x-axis represents sRNA length (nt), and the y-axis shows RNA abundance in reads per million (RPM). Stages include In Planta Appressorium (PA), Biotrophic Phase (BP), Necrotrophic Phase (NP), In Vitro Appressorium (VA), and Mycelia (MY). Three biological replicates are represented by distinct colors: blue, mustard-yellow, and orange.

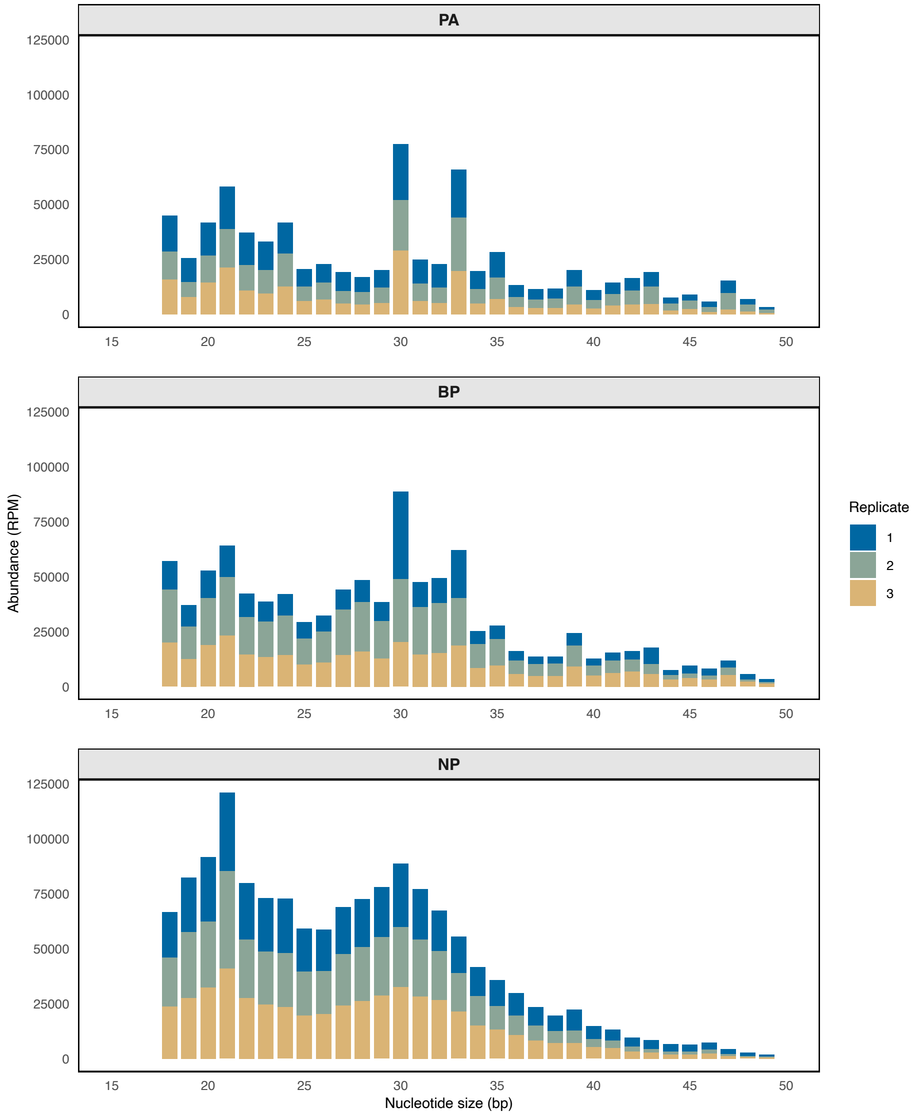


**Figure S7. Size distribution of gene-derived sRNAs in *A. thaliana* during infection by *C. higginsianum*.**
The x-axis represents sRNA length (nt), and the y-axis shows RNA abundance in reads per million (RPM). Infection stages include In Planta Appressorium (PA), Biotrophic Phase (BP), and Necrotrophic Phase (NP). Three biological replicates are represented by distinct colors: blue, green, and mustard-yellow.

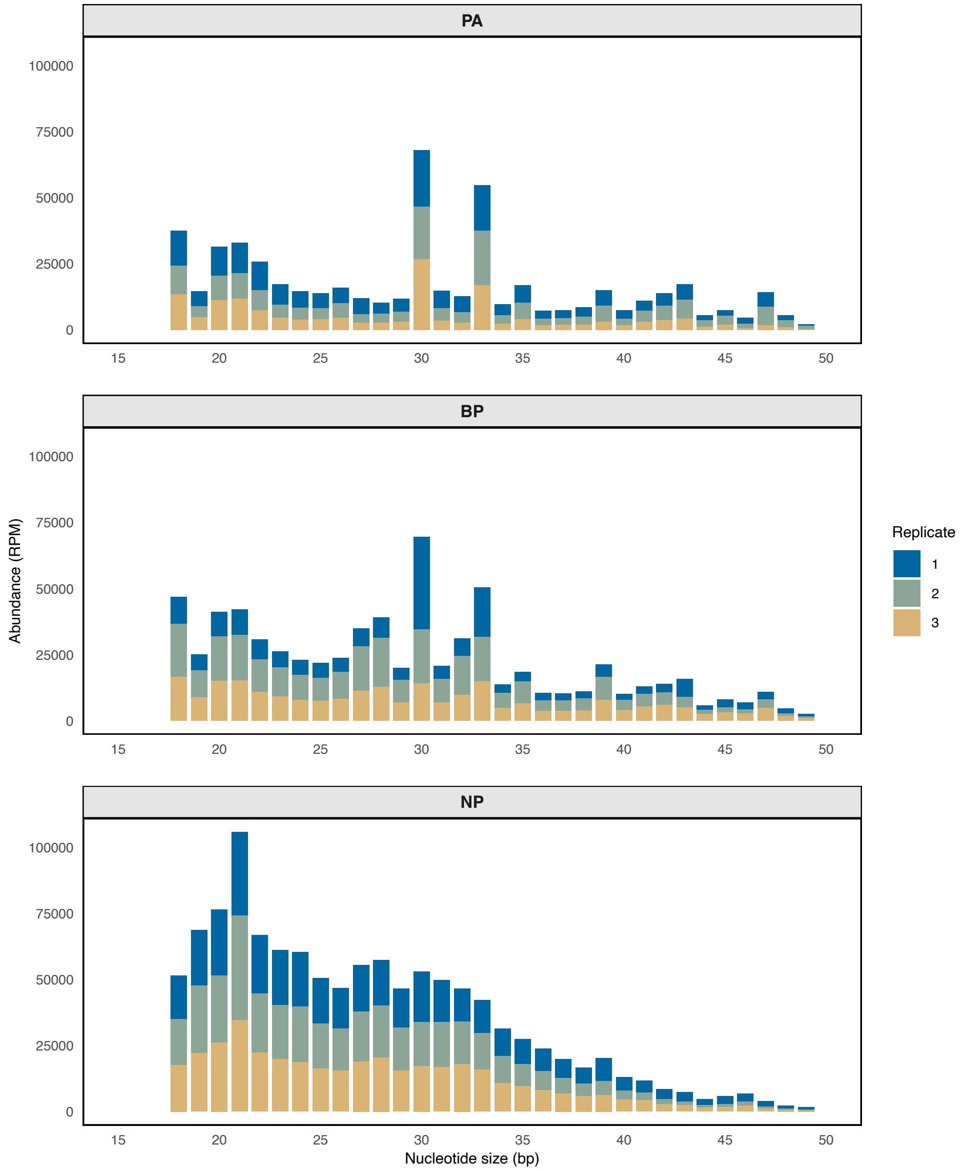


**Figure S8. Size distribution of rRNA-derived sRNAs in *A. thaliana* during infection by *C. higginsianum*.**
The x-axis represents sRNA length (nt), and the y-axis shows RNA abundance in reads per million (RPM). Infection stages include In Planta Appressorium (PA), Biotrophic Phase (BP), and Necrotrophic Phase (NP). Three biological replicates are represented by distinct colors: blue, green, and mustard-yellow.

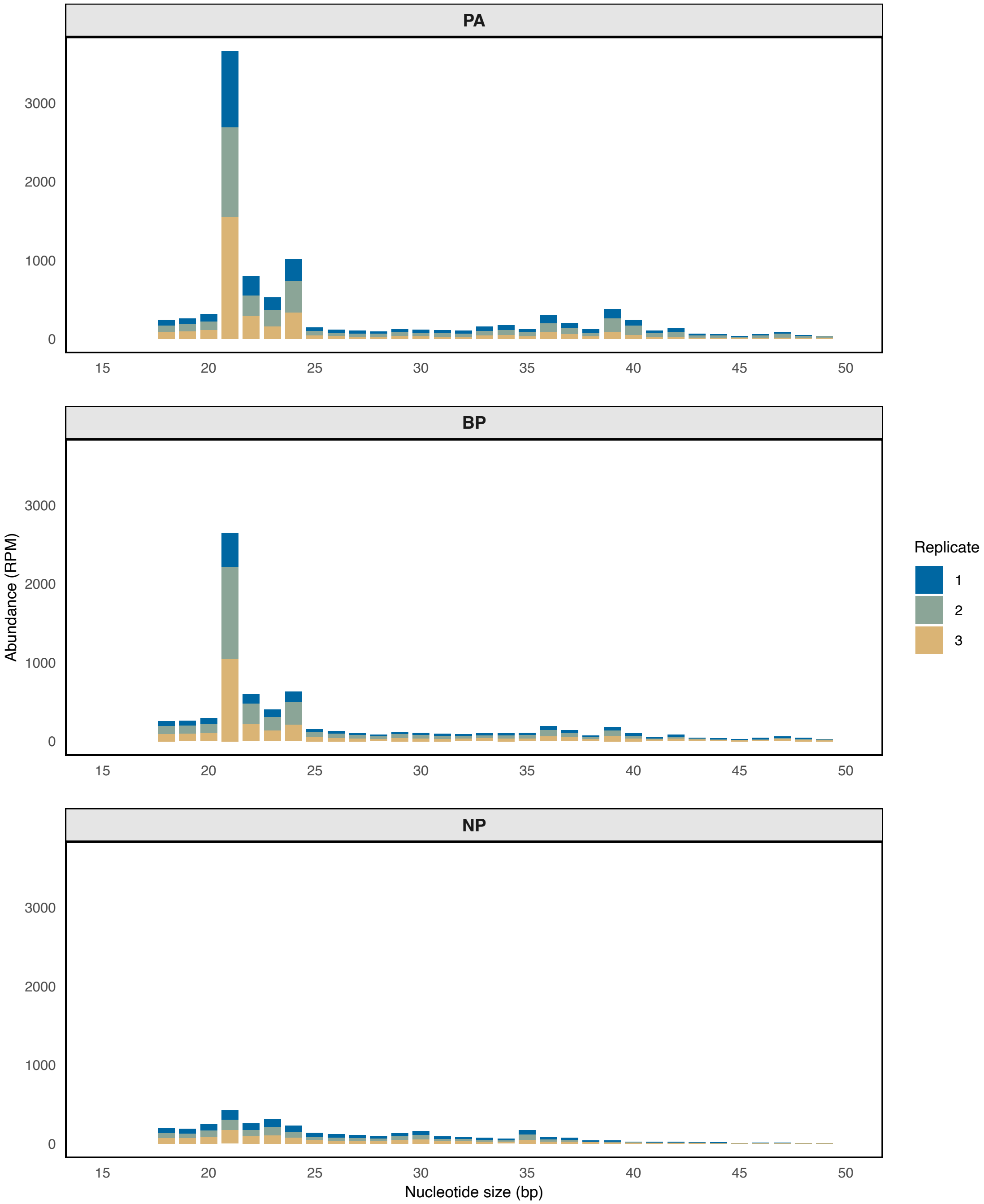


**Figure S9. Size distribution of ncRNA-derived sRNAs in *A. thaliana* during infection by *C. higginsianum*.**

The x-axis represents sRNA length (nt), and the y-axis shows RNA abundance in reads per million (RPM). Infection stages include In Planta Appressorium (PA), Biotrophic Phase (BP), and Necrotrophic Phase (NP). Three biological replicates are represented by distinct colors: blue, green, and mustard-yellow.

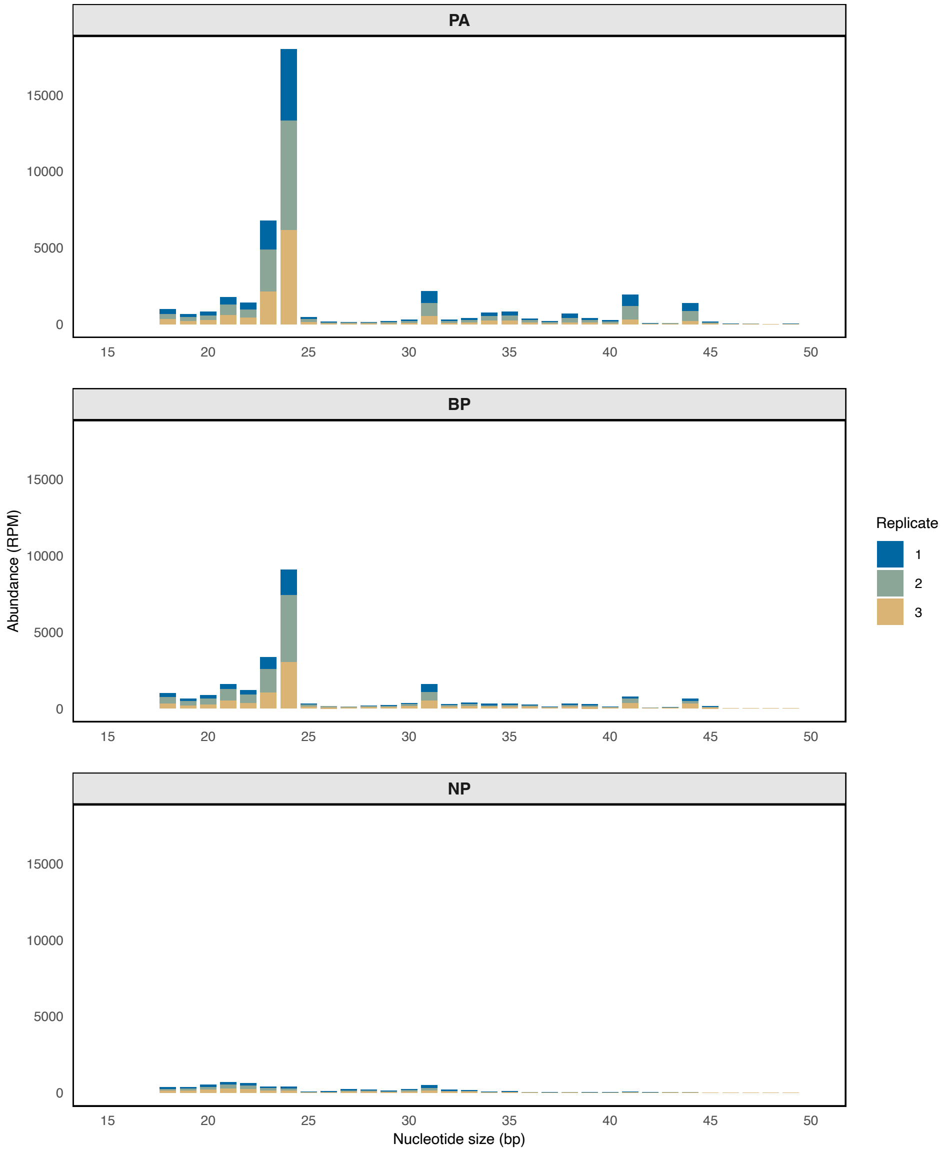


**Figure S10. Size distribution of TE-derived sRNAs in *A. thaliana* during infection by *C. higginsianum*.**
The x-axis represents sRNA length (nt), and the y-axis shows RNA abundance in reads per million (RPM). Infection stages include In Planta Appressorium (PA), Biotrophic Phase (BP), and Necrotrophic Phase (NP). Three biological replicates are represented by distinct colors: blue, green, and mustard-yellow.

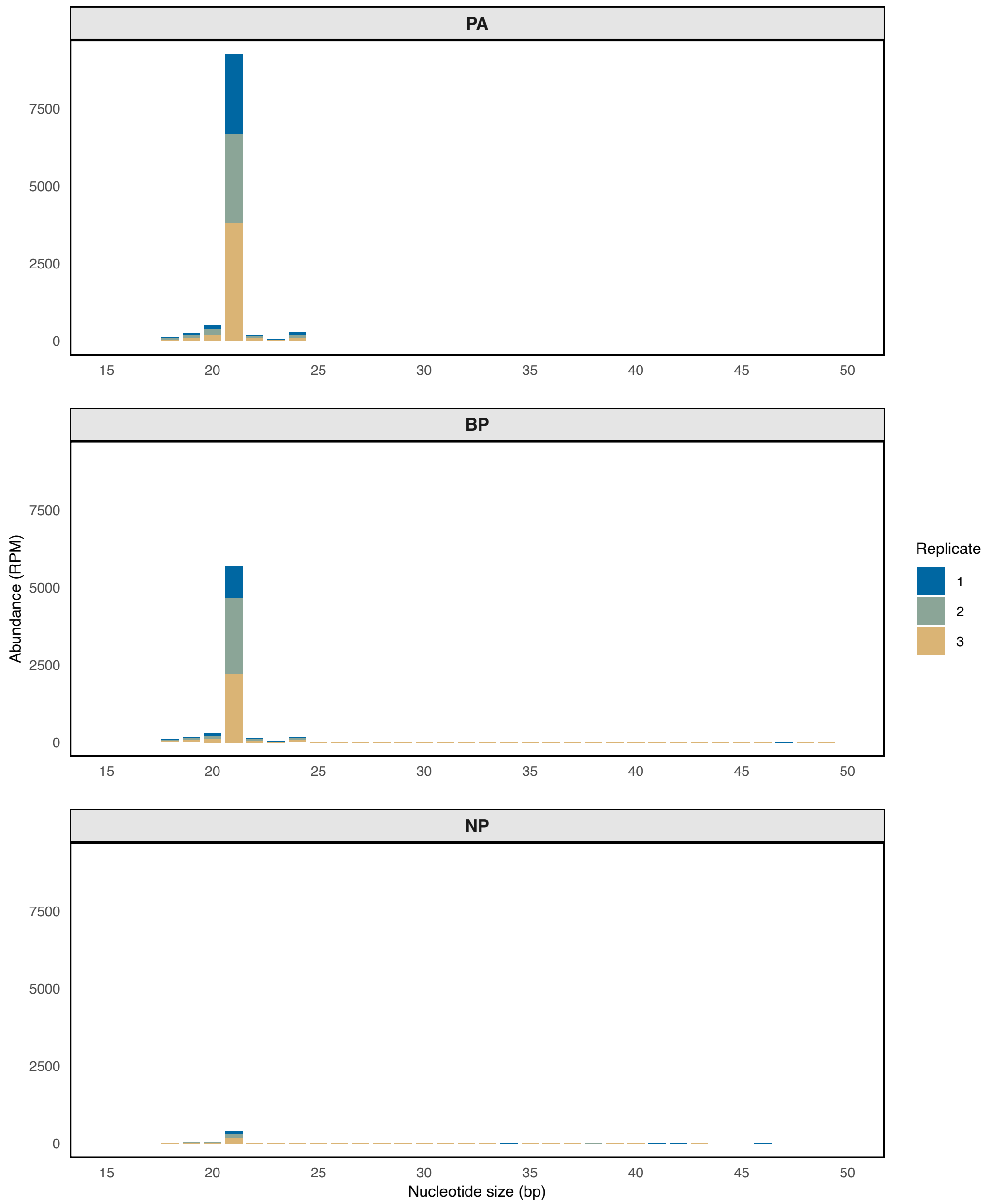


**Figure S11. Size distribution of miRNA-derived sRNAs in *A. thaliana* during infection by *C. higginsianum*.**
The x-axis represents sRNA length (nt), and the y-axis shows RNA abundance in reads per million (RPM). Infection stages include In Planta Appressorium (PA), Biotrophic Phase (BP), and Necrotrophic Phase (NP). Three biological replicates are represented by distinct colors: blue, green, and mustard-yellow.

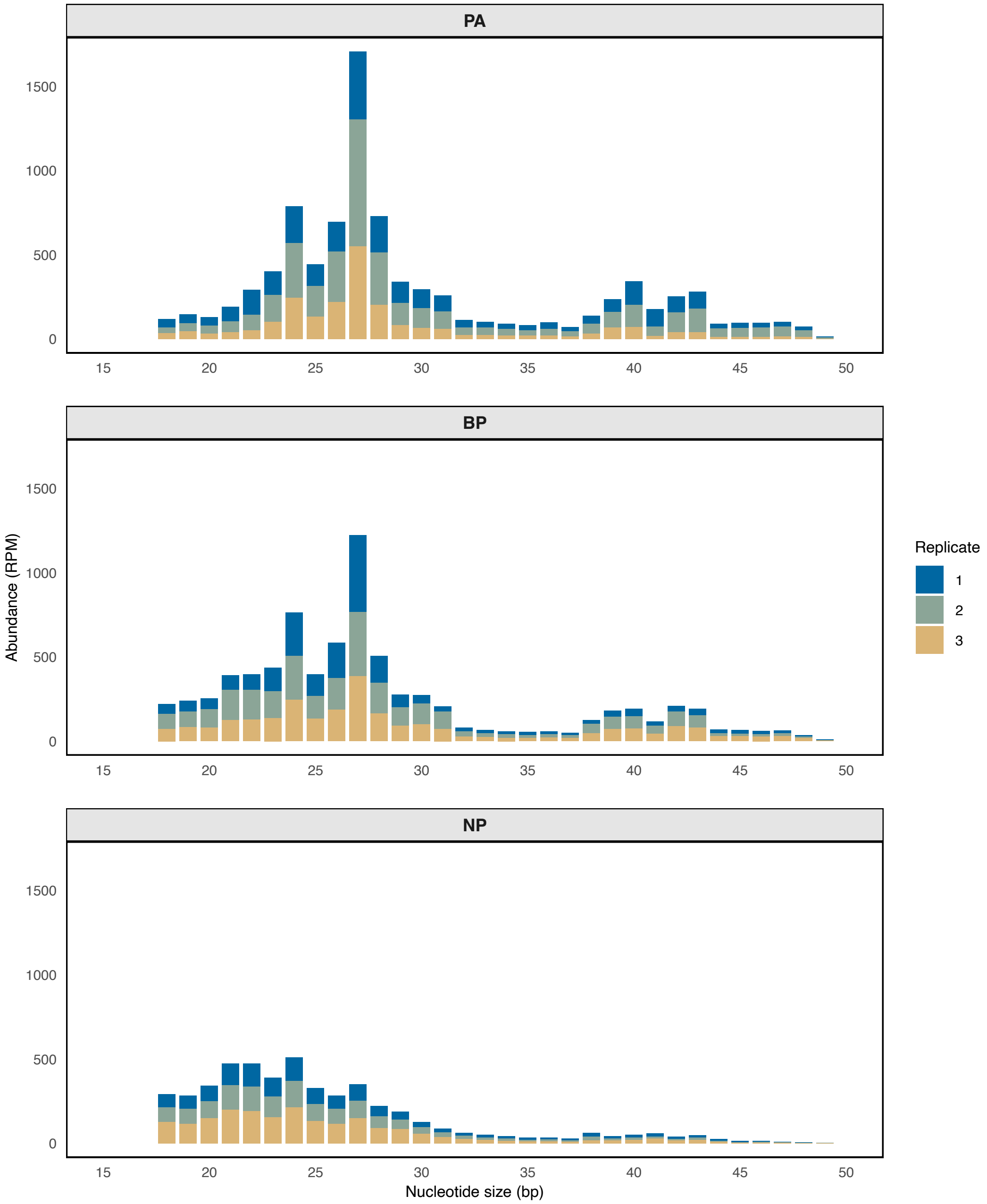


**Figure S12. Size distribution of snRNA-derived sRNAs in *A. thaliana* during infection by *C. higginsianum*.**
The x-axis represents sRNA length (nt), and the y-axis shows RNA abundance in reads per million (RPM). Infection stages include In Planta Appressorium (PA), Biotrophic Phase (BP), and Necrotrophic Phase (NP). Three biological replicates are represented by distinct colors: blue, green, and mustard-yellow.

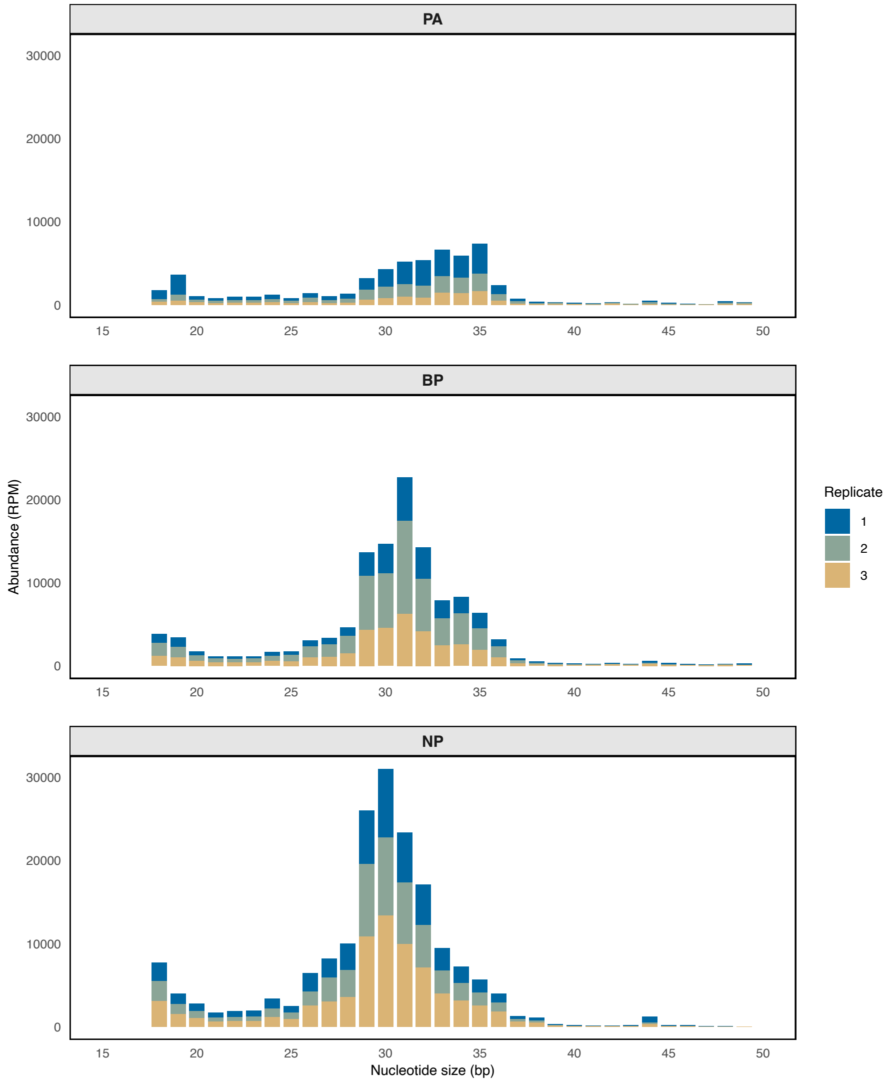


**Figure S13. Size distribution of tRNA-derived sRNAs in *A. thaliana* during infection by *C. higginsianum*.**
The x-axis represents sRNA length (nt), and the y-axis shows RNA abundance in reads per million (RPM). Infection stages include In Planta Appressorium (PA), Biotrophic Phase (BP), and Necrotrophic Phase (NP). Three biological replicates are represented by distinct colors: blue, green, and mustard-yellow.

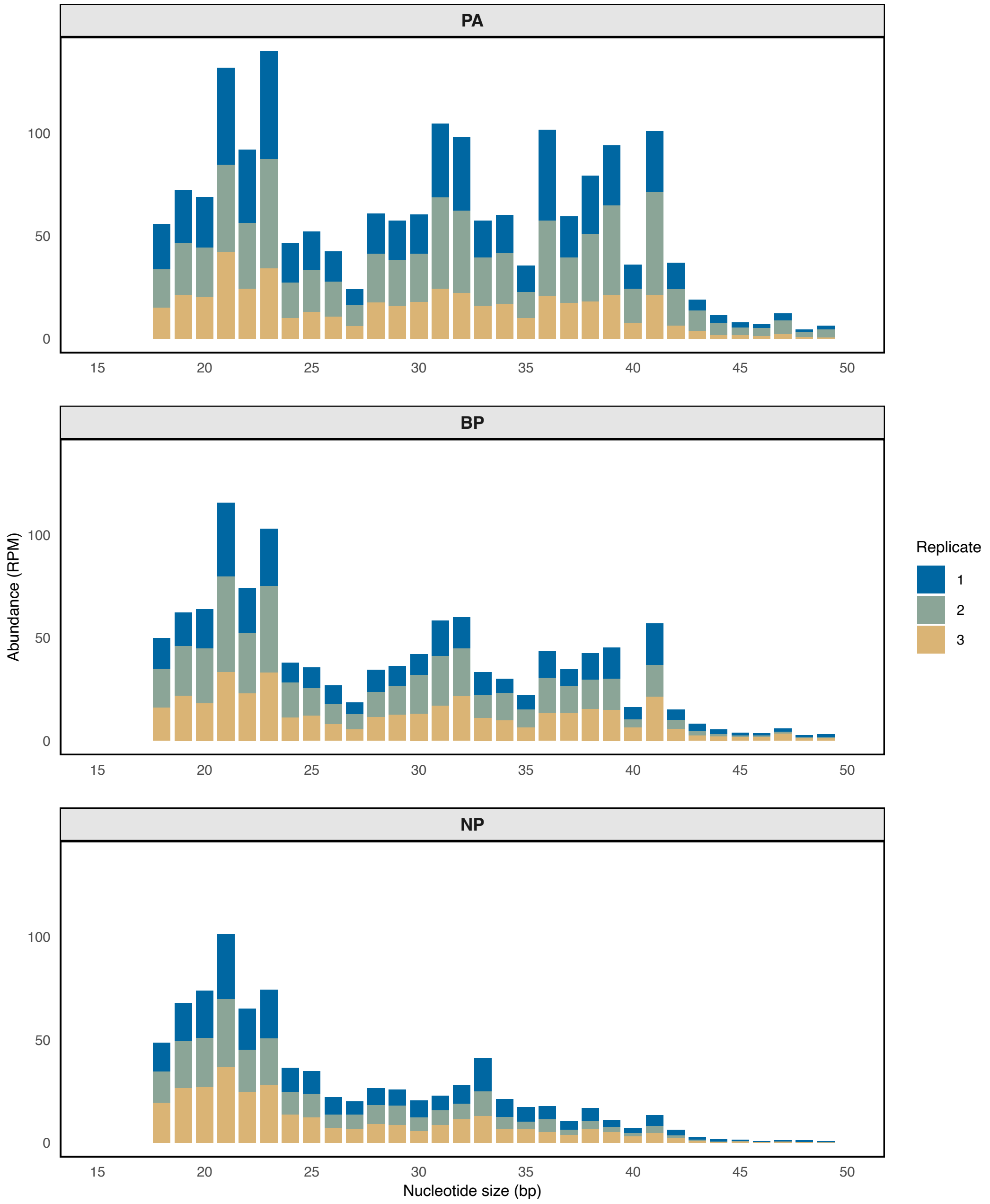


**Figure S14. Size distribution of snoRNA-derived sRNAs in *A. thaliana* during infection by *C. higginsianum*.**

The x-axis represents sRNA length (nt), and the y-axis shows RNA abundance in reads per million (RPM). Infection stages include In Planta Appressorium (PA), Biotrophic Phase (BP), and Necrotrophic Phase (NP). Three biological replicates are represented by distinct colors: blue, green, and mustard-yellow.
